# Metabolomic and lipidomic shifts underpin physiological acclimation to thermal stress in the European green crab (*Carcinus maenas*)

**DOI:** 10.64898/2026.05.08.723818

**Authors:** Yaamini R. Venkataraman, Sara K. Shapiro, Mikayla Newbrey, Carolyn K. Tepolt

## Abstract

Many marine invertebrates are characterized by broad and highly plastic thermal limits, though the dynamic molecular mechanisms that enable extended thermal acclimation remain poorly understood. A classic example is the green crab (*Carcinus maenas*), which is a prolific and damaging non-indigenous species. Using a 22-day thermal exposure to cold (5°C), ambient (13°C), or warm (30°C) temperatures, we characterized plastic shifts in *C. maenas* performance using respirometry and time-to-right. We then used untargeted metabolomics and lipidomics analysis of heart tissues from days 4 and 22 to identify the molecular mechanisms underpinning plastic responses over time. Crabs at 30°C exhibited higher oxygen consumption rates than counterparts at 5°C. Interestingly, oxygen consumption rate increased over time at both temperatures, indicating thermal plasticity of aerobic respiration. Temperature-dependent metabolic reprogramming was employed by crabs to sustain aerobic respiration across temperature. Catabolism of branched-chain amino acids was important for energy production at elevated temperatures, while catabolism of arginine may have sustained the minimal energy needs of crabs exhibiting metabolic depression at cold temperatures. Righting response was positively correlated with temperature, and did not exhibit any changes over time. Lipidome remodeling consistent with homeoviscous adaptation could have enabled motor activity across temperature. Higher abundances of saturated and monounsaturated lipids likely provided structural integrity to cell membranes at 30°C, while lower abundances of these compounds may have enabled membrane fluidity at 5°C. Our work demonstrates the importance of ongoing molecular reprogramming in long-term acclimation, even when whole-animal physiology remains relatively stable.

**Summary Statement:** This study demonstrates how the highly invasive green crab regulates metabolite and lipid pathways over time to maintain physiological performance across different temperatures.

## Introduction

Temperature fundamentally shapes organismal physiology, particularly in ectotherms (Somero et al., 2017). However, it can be difficult to tease apart the various impacts of temperature on physiology across time and biological hierarchy (Schulte, 2015; Schulte et al., 2011). Thermal exposure duration can elicit time-dependent metabolic and acclimation responses (Angilletta, 2009; Schulte et al., 2011) that are seemingly contradictory when comparing whole-organism responses with physiological and molecular biomarkers (Zou et al., 2021; Zou et al., 2023). Comparing acute (a few hours) and chronic (days-weeks) thermal responses can also highlight the induction and maintenance of an acclimation response (Evensen et al., 2021), or the waning of that same response. Experiments that examine both short- and long-term thermal response across the whole organism, physiological pathway, and molecular levels are necessary for a comprehensive understanding of thermal limits and how they change over time.

Marine invertebrates, especially those found in the intertidal zone, have evolved complex mechanisms with which to deal with the thermal stress they regularly experience. Intertidal temperatures change rapidly and dramatically across daily and seasonal cycles, providing the opportunity to study mechanisms underlying phenotypic plasticity at a range of time scales (Denny et al., 2011; Helmuth and Hofmann, 2001; Leeuwis and Gamperl, 2022). Prior work has shown consistent changes in metabolic markers with cyclical changes in environmental conditions in marine invertebrates such as oysters (Hamdoun et al., 2003) and barnacles (Broitman et al., 2021). These molecular changes can mediate changes to whole-organism physiology that improve thermal tolerance (Chen et al., 2018; Kelley et al., 2011). Although intertidal species can respond metabolically to predictable cycles of tide, day, and season, or to acute temperature shifts (Madeira et al., 2018), they may be living closer to their thermal limits with little ability to mount phenotypically plastic responses long-term (Harley et al., 2006). Additional long-term studies with intertidal organisms are necessary for interrogating physiological responses to environmental conditions and investigating possible tradeoffs between thermal tolerance and plasticity (Barley et al., 2021).

The integration of whole-organism physiology with untargeted molecular methods, particularly metabolomics and lipidomics, can elucidate the mechanisms used by organisms like marine invertebrates to withstand chronic thermal stress (Venkataraman and Huffmyer, 2025). Metabolomics, or the study of direct and indirect products of metabolic pathways, can provide insight into energetic pathways used to maintain homeostasis in response to a stressor (Johnson et al., 2016; Lankadurai et al., 2013; Liu and Locasale, 2017; Venkataraman and Huffmyer, 2025). Lipidomics facilitates the study of fatty acid and lipid content, both of which are important for short- and long-term energy storage as well as homeoviscous adaptation (Rey et al., 2022; Venkataraman and Huffmyer, 2025; Wenk, 2005). The connection of both of these techniques to energy metabolism allows researchers to interrogate an organism’s metabolic plasticity (Fendt et al., 2020; Jia et al., 2019). For example, northern shrimp (*Pandalus borealis*) exhibit metabolome (Guscelli et al., 2023) and lipidome (Feugere et al., 2025) reprogramming in response to acidification and temperature stress. A similar integrative approach revealed that Dungeness crab (*Cancer magister*) responds to combined ocean acidification and hypoxia by reducing the activity of ATP-consuming processes (Wanamaker et al., 2019). Pairing these molecular methods with physiological assays can provide insight into mechanistic responses that may not be apparent at the whole-organism level (Venkataraman and Huffmyer, 2025). Coral larvae exhibited metabolic reprogramming when exposed to elevated temperature without a concurrent decrease in survival (Huffmyer et al., 2024). Similarly, exposure of American lobsters (*Homarus americanus*) to ocean acidification resulted in broad metabolic reprogramming that was not associated with changes to resting metabolism, suggesting beneficial plasticity of the metabolic response (Noisette et al., 2021). Employing these tools can elucidate mechanisms of susceptibility and resilience as organisms respond to environmental stressors.

The European green crab (*Carcinus maenas*) is an ideal candidate for using multi-faceted thermal experiments to understand how thermal limits are shaped and reshaped physiologically. Green crabs are a prolific non-indigenous species in North America, characterized by broad thermal tolerance and exceptionally flexible thermal physiology (Tepolt, 2024). Acute thermal ramp studies have identified CT_max_ above 35°C for righting response cessation (Kelley et al., 2011), heart rate breakpoint (Tepolt and Somero, 2014), and mortality (Cuculescu et al., 1998), and cardiac activity is maintained down to at least -0.8 °C (Frederich et al., 2000; Tepolt and Somero, 2014). Like many intertidal invertebrates, *C. maenas* exhibits thermal plasticity that is responsive to natural environmental variation including tidal cycles (Nancollas and McGaw, 2021) and seasons (Cuculescu et al., 1998; Hopkin et al., 2006). The maintenance of broad thermal tolerance and extensive plasticity likely reflects several complementary mechanisms: “dumping” hemocyanin-bound oxygen to extend heat tolerance (Giomi and Pörtner, 2013; Jost et al., 2012); increasing mitochondrial density and capacity (McGaw and Whiteley, 2024); reducing hemolymph magnesium to support movement at cold temperatures (Frederich et al., 2000; Wittmann et al., 2010); and altering membrane lipid composition to maintain fluidity and permeability across a range of temperatures (Chapelle, 1978; Chapelle, 1986; El Babili et al., 1997). Previous research has also investigated the association of specific biomarkers with response to thermal stress in *C. maenas*, and found that there was no difference in HSP 70 in *C. maenas* across a 24-35.1°C or 12-34°C temperature range (Jost et al., 2012; Madeira et al., 2014), and no change in AMP-activated protein kinase activity across the same 12-34°C range (Jost et al., 2012). While Jost et al. (2012) did identify an increase in lactate abundance at 34°C, suggesting that crabs employ anaerobic metabolism between 32-34°C, the limited utility of specific biomarkers suggests that untargeted assays may better illuminate the pathway-level changes that underlie the sudden onset of a thermal stress response in this species.

In this study, we used an integrative approach to examine *C. maenas* response to chronic thermal stress. Crabs were exposed to either warm (30°C) or cold (5°C) temperature conditions for 22 days, and compared with a non-stressful 13°C control treatment. These temperatures were selected based on previous work (Tepolt and Somero, 2014) with the goal of inducing the metabolic reprogramming expected in stressful thermal conditions. Throughout the experiment, we measured survival and performance including righting response and oxygen consumption rate. We expected that crabs would initially maintain performance at thermal extremes, but that compensation would wane as exposure continued. We also used untargeted metabolomics and lipidomics to investigate the molecular mechanisms activated by *C. maenas* at the beginning and end of the exposure. We expected that metabolomes and lipidomes profiled earlier in the exposure period would highlight pathways utilized to maintain performance, while pathways identified in samples taken at the end of the exposure would be indicative of a stress response. Our work demonstrates how metabolomic and lipidomic reprogramming underpins shifts in whole-organism physiology, and how these impacts are dependent on time.

## Materials and Methods

### Experimental Design

Green crabs were obtained on June 29, 2022 from a bait shop in Bourne, MA, USA, that collected crabs locally. Crabs with orange or red integuments were excluded, as these colors indicate animals that have spent an extended period in intermolt and may have different environmental tolerances than those with greener integument colors (Reid et al., 1997). Selected crabs were placed in 21 cm x 26 cm x 41 cm (22.386 L) tanks, with 14-15 crabs in each of six tanks filled with 50 micron filtered raw seawater from Vineyard Sound, MA, USA. To maintain water temperature at ∼13°C in a static system, tanks were placed in a flowing water bath. Two HOBO Data Loggers (Onset, USA) — one at the water surface and one at the bottom of the tank — were used to monitor tank temperature every 15 minutes. An air stone (Quickun Aquarium Air Stone) attached to a 10L aerator (Whisper® Aquarium Air Pumps, Tetra®) was included in each tank. An aquarium filter (MARINA i25) was used to maintain water quality in each tank, with filter cartridges replaced every five days throughout the course of the acclimation period and experiment. The tanks were kept on a 12-hour light, 12-hour dark cycle (Tepolt and Somero, 2014).

All crabs were acclimated at 13°C for seven days, then moved to experimental temperatures for 22 days (Figure 1). Crabs from each acclimation tank were shuffled between experimental tanks to ensure an even distribution of sex, integument color, and size in each experimental tank. Two tanks were maintained at 13°C throughout the experiment as controls. For the elevated temperature treatment, two 300-Watt Deluxe Titanium Heating Tubes (Finnex, USA) with external 1650-Watt temperature controllers (bayite, China) were placed in each experimental tank to maintain a 30°C treatment. The two remaining tanks were moved into a cold room set at 5°C to achieve a 5°C cold treatment. Data from replicate HOBO loggers in each tank were used with a one-way Analysis of Variance (ANOVA; `aov` from `stats` package from R Statistical Programming v 4.5.1 (R Core Team, 2024)) to confirm that temperature treatments were statistically different from each other.

**Figure 1.**
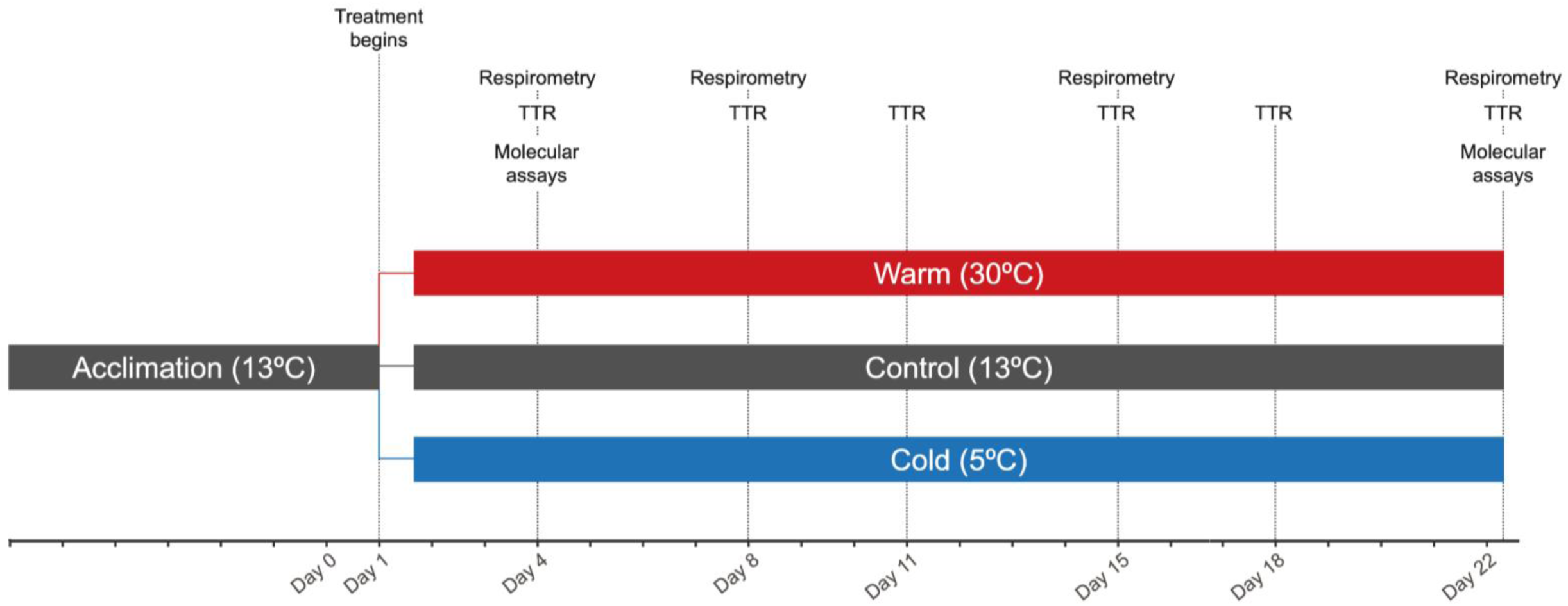
Experimental timeline, including sampling points for time-to-right (TTR), respirometry, and molecular assays (metabolomics and lipidomics). Mortality was checked daily starting on day 0, and demographic variables (carapace width, crab weight, integument color, and sex) were assessed on day 1 and after all time-to-right assays. Heart tissue was sampled for metabolomics and lipidomics analyses.

Throughout the acclimation period and the experiment, tanks were checked daily for mortality. Ammonia concentrations were evaluated every other day using an API® Ammonia Test Kit. Particulate waste was removed daily. Sixty percent of water was replaced every three days and when ammonia concentrations were above 1.0 ppm. During these water changes, ½ tsp of conditioner (Kordon Amquel Plus Aquarium Water Conditioner) was added to tanks when ammonia exceeded this threshold. All crabs were fed daily, with crabs in control (13°C) and elevated (∼30°C) temperature tanks given ½ teaspoon of food (Crab Cuisine, Hikari USA Inc.) and crabs in cold (∼5°C) temperature tanks given ¼ teaspoon of food since they consumed less. Food quantity was sufficient to satiate crabs as there was always a small amount of uneaten food the next day.

Crab demographic data was compared to ensure these parameters were consistent between temperature treatments. Carapace width (mm), crab weight (g), integument color, and sex were compared between temperature treatments at day 1 and 22. Kruskal-Wallis tests were used to compare mean carapace width and weight, and chi-squared tests were used to assess differences in integument color and sex distributions across the different temperatures (`kruskal.test` from R `stats` package v.4.5.1).

### Performance Assays

#### Survivorship

Survivorship was assessed using the R packages `survival` v.3.5.5 (Therneau, 2023; Therneau and Grambsch, 2001) and `ggsurvfit` v.0.3.0 (Sjoberg D, Baillie M, Haesendonckx S, Treis T, 2023). Data were formatted with `Surv`, with right-censoring to account for *C. maenas* that survived until the end of the experiment. Overall differences in *C. maenas* survival due to temperature treatment were tested with a non-parametric Kaplan-Meier survival analysis (`survdiff`), and Kaplan-Meier curves were plotted (`survfit2`). A Cox proportional hazards regression model (`coxph`) was then used to calculate the change in chance of death for the 30°C or 5°C treatment compared to the 13°C control.

#### Time-to-right

Time-to-right (TTR) was used to compare the whole-animal physiological performance of *C. maenas* between temperature treatments on days 4, 8, 11, 15, 18, and 22 (**Figure 1**). TTR was defined as the the number of seconds *C. maenas* spent on righting behavior (defined in (Young et al., 2006)), as well as the “hesitation time” before moving the fifth pereopod pair after crabs are placed dorsal side up. Seven crabs in each tank were randomly selected for each TTR assay at their experimental temperature. Crabs were placed upside-down, with the dorsal surface on the bottom of the tank. If a crab took longer than 90 seconds to right, crabs were manually righted and recorded as “not righting” for that trial. Crabs were allowed to rest 10 seconds, then the assay was repeated twice for a total of three TTR measurements per crab. Additional trials were not added for each time the crab did not right itself. At the end of the experiment, TTR was tested for all surviving crabs. All crabs were weighed, measured, sexed, and assessed for integument color and missing legs after completing the TTR trials. Since righting is initiated by moving the fifth pair of pereopods (Young et al., 2006), crabs missing both fifth pereopods were excluded from this assay. Prior to analysis, replicate TTR measurements for each individual crab were averaged, excluding trials where the crab did not right itself, and log-transformed.

A generalized linear model (GLM; `glm` from R `stats` package (R Core Team, 2024)) was used to assess differences in averaged TTR due to the influence of treatment, experimental day, the interaction between treatment and day, integument color, carapace width, weight, and whether or not the crab was missing legs. A backwards deletion approach was used for model construction (`step` from R `stats` package (R Core Team, 2024)), with the best fit model having the lowest Akaike information criterion (AIC) score. Factors were considered significant when *P*-value < 0.05. Normality and homoscedasticity were verified visually. Pairwise tests were conducted with `emmeans` v.1.11.0 for any significant factor (Lenth et al., 2018). Pairwise comparisons were considered significant when adjusted *P*-value < 0.05.

#### Oxygen consumption

Rate of oxygen consumption was used to measure metabolic performance of *C. maenas* at the crabs’ treatment temperatures on days 4, 8, 15, and 22 (**Figure 1**). Crabs (n = 3 per tank) from the TTR trial that weighed roughly 20 g were placed in 90 mm x 50 mm (270 mL) glass respirometry chambers with FireSting oxygen sensor spots (PS-OXSP5; PyroScience, Germany) and covered with dark material to prevent light from entering. These containers were covered with mesh to allow water exchange, and placed back into an aerated tank at the crab’s treatment temperature. After the one hour of acclimation, respirometry chambers were refilled with aerated water at the treatment temperature, then sealed with parafilm and a glass lid. The respirometry chambers were placed in a water bath at the treatment temperature, and weights were placed on top of the lid to further prevent oxygen intrusion. Percent oxygen saturation was measured in one-second intervals until it reached 75% saturation. The crabs were then removed from respirometry chambers, and the chambers were quickly resealed to measure the background oxygen consumption for at least twenty minutes. Salinity measurements were obtained for each experimental tank using a refractometer. Prior to each assay, respirometry chambers were cleaned with 70% ethanol, and bare optical fibers for FireSting devices (PS-SPFIB-BARE) were calibrated using ambient air as the upper limit.

Oxygen consumption rate was compared between temperature treatments and timepoints. First, rates were calculated for individual respirometry chambers. Once data were cleaned, dissolved oxygen (µmol/L) was calculated using percent air saturation, temperature (°C), and pressure (mbar) information from the individual chambers and a salinity measurement obtained from each tank with the `conv_o2` function from the R package `respirometry` v.2.0.0 (Birk, 2023). For time points between 80-100% air saturation, linear models were fitted to oxygen consumption data to obtain regression slopes for dissolved oxygen (`fit` from the R package `generics` v.0.1.3 (Wickham et al., 2022)) and adjusted R-squared values (`glance` from `generics` package). Next, a “blank” oxygen consumption rate was calculated by fitting linear models to the last ten minutes of background oxygen consumption measurements. The background slope was subtracted from the crab oxygen consumption slope to obtain a blank-corrected slope. A GLM was used to evaluate treatment, day, and the interaction between treatment and day on log-transformed blank-corrected oxygen consumption rates. Crab sex, integument color, carapace width, weight, and whether or not the crab was missing legs were also used as explanatory variables. A backwards deletion approach (`step`) was used to find the most parsimonious model with the lowest AIC score. Factors were considered significant when *P*-value < 0.05. Normality and homoscedasticity were verified visually. Pairwise tests were conducted with `emmeans` v.1.11.0 for any significant factor (Lenth et al., 2018). Pairwise comparisons were considered significant when adjusted *P*-value < 0.05.

### Molecular Assays

#### Spectra acquisition and quantification

Crab heart tissue was collected on day 4 (n = 6 per temperature treatment) and day 22 (n = 21 for 13°C, n = 21 for 5°C, and n = 10 for 30°C) for molecular assays (**Figure 1**). After live dissection by severing the central ganglion, heart tissue was flash frozen on dry ice and stored at -80°C. Samples were sent to the West Coast Metabolomics Center, Davis, CA for sample preparation and untargeted analysis of metabolites and lipids. Metabolites were isolated following methods in Phillips et al. (2024) (additional information available in **Appendix 1**). Processed samples underwent Automated Liner Exchange-Cold Injection System (Gerstel Corporation) Gas Chromatography and Time-of-Flight Mass Spectrometry (ALEX-CIS GCTOF MS) using a Leco Pegasus IV mass spectrometer for positively and negatively charged metabolites following parameters from Fiehn et al. (2008). Resultant spectra were processed using the UC Davis BinBase workflow (Fiehn et al., 2005). The sum of all spectral peak heights was calculated for identified metabolites within each sample (metabolite total ion chromatogram, “mTIC”), and individual spectra were then normalized by the average mTIC of all samples.

Lipids were extracted from heart tissue following methods in Matyash et al. (2008) modified so lipid extracts were on the top layer during phase separations to ensure lipids were not contaminated by proteins or polar compounds. Lipidomics samples underwent Electrospray Ionization Quadrupole Time-of-Flight Tandem Mass Spectrometry (ESI QTOF MS/MS). Mass spectrometry was separated into a negative mode and a positive mode to capture compounds of varying charges. Lipid spectra were processed using MS-DIAL version 4.92 (Tsugawa et al., 2015) and cleaned with MS-FLO (DeFelice et al., 2017), followed by annotation using the Fiehn laboratory LipidBlast spectral library (Kind et al., 2013) and manual scans of the data, and peak verification with MassHunt Quant scanner. Data were normalized using the total sum of the internal standards.

#### Analysis of known compounds

Metabolomic and lipidomic data were analyzed separately using similar workflows following best practices outlined in Venkataraman and Huffmyer (2025). Broadly, 1) an unsupervised Permutational Multivariate Analysis of Variance (perMANOVA) was used to examine treatment effects on compound abundance; 2) a supervised Partial Least Squares Discriminant Analysis (PLS-DA) was used to identify specific compounds contributing to statistically significant variation between treatment groups; and 3) an ANOVA-Simultaneous Component Analysis (ASCA) was used to classify compounds into groups based on similar abundance patterns across temperature and/or day.

Overall abundance patterns of all detected compounds were first assessed visually using a Principal Components Analysis (PCA). Missing data were replaced with a zero, and data were scaled for the PCA. Significant influences of temperature, day, and their interaction were quantified using a global perMANOVA using a euclidean distance matrix created with scaled data (`adonis` from the R package `vegan` v.2.6-6.1 (Oksanen et al., 2024)). A beta dispersion test (`betadisper` from the R package `vegan` v.2.6-6.1) was used to determine if any significant effects detected in the perMANOVA were solely due to centroid differences or combined centroid and variance differences. The PCA, perMANOVA, and beta dispersion tests were repeated on the subset of compounds that were identified by the sequencer (“known compounds”) to confirm the overall patterns did not change between the full data set and known subset.

Significant effects from the perMANOVA were used in a subsequent supervised PLS-DA (`plsda` from the R package `mixOmics` v.6.28.0 (Rohart et al., 2017)) with log-transformed abundance data from known compounds. The number of components and features used in the PLS-DA were validated using permutation tests. An Mfold cross validation (`perf`) with six folds (`folds = 6`) was repeated 50 times (`nrepeat = 50`) to determine the number of components to use for the PLSDA. The optimal number of features (known metabolites or lipids) was determined using M-fold validation on a series of sparse PLS-DAs (`tune.splsda.srbct`) with six folds (`folds = 6`), 50 repeats (`nrepeat = 50`) and the number of components determined in the previous validation step (`ncomp`). The influence of specific compounds in differentiating pairs of temperature x day contrasts was assessed by obtaining variable importance projection (VIP) scores using the `PLSDA.VIP` function within the `mixOmics` package. Higher VIP scores indicate an individual compound’s abundance is more influential in driving differences between different groups assessed by the PLS-DA model. Significant compounds from pairwise contrasts between temperatures and/or days (“pairwise VIP”) were identified by comparing log-normalized compound abundance (`t.test` from R `stats` package (R Core Team, 2024)). Compounds were considered pairwise VIP if their VIP score ≥ 1 and their adjusted t-test *P*-value < 0.05.

After determining how temperature, day, and/or their interaction shaped overall abundance patterns, an ANOVA-Simultaneous Component Analysis (ASCA) was used to identify groups of compounds with similar abundance patterns in various conditions (Bertinetto et al., 2020). This method is useful for applying multivariate methods to analyzing datasets with more variables than observations (Bertinetto et al., 2020). A model was constructed using the `ALASCA` (v.1.0.17) package in R (Jarmund et al., 2022) to analyze log-transformed compound abundance as a function of fixed (significant effects from PCA) and random (individual crab) effects. The model was validated with bootstrapping with replacement for 100 iterations (Pattengale et al., 2010). Data for PCs that explained > 10% of the variance were visualized using effect plots (`plot(type = “effect”)`). Compounds could be positively loaded onto the PC (have the same abundance patterns as visualized in the effect plot) or negatively loaded onto the PC (have opposite abundance patterns than what is visualized in the effect plot). Information for the 20 compounds with the highest PC scores were also extracted for each PC (`get_loadings(limit = 10)`) to assess if any pairwise VIP were strongly correlated with any PC.

#### Functional analysis

Functional analysis was conducted using two methods for each of the metabolomics and lipidomics data sets. Enrichment analysis was first conducted for each set of pairwise VIP compounds. For pairwise VIP metabolites, combined enrichment and pathway topology analysis was conducted using the Pathway Analysis module in the MetaboAnalyst v.6.0 interface (Pang et al., 2024). Kyoto Encyclopedia of Genes and Genomes (KEGG) pathways associated with *Drosophila melanogaster* were used as a reference. Pathways were considered significantly enriched if FDR < 0.05. For pairwise VIP lipids, target enrichment analysis was conducted with LION/web (Molenaar et al., 2019), a web-based GUI that performs enrichment analysis on lipid-specific ontology terms (i.e. LION terms) using a one-tailed Fisher’s exact test. LION terms were considered significantly enriched if FDR < 0.05. Similar enrichment methods were used to understand overrepresented pathways in ASCA-derived PCs that explained > 10% of total variance. The 20 compounds with the highest PC scores were used for enrichment.

The function of pairwise VIP compounds was also assessed with respect to compound classes of interest. Temperature is known to influence energy metabolism (Pörtner, 2010; Sokolova et al., 2012; Weihrauch and McGaw, 2024), and cold temperatures are specifically associated with magnesium-blockage of calcium channels in *C. maenas* (Frederich et al., 2000; Wittmann et al., 2010). Therefore, the involvement of pairwise metabolites was assessed in aerobic respiration (glycolysis, glycogenolysis, citric acid cycle) pathways, anaerobic respiration (glycolysis, glycogenolysis, and lactic acid fermentation) pathways, and calcium signaling pathways. Changes in abundance were examined for pairwise VIP lipids that were part of the triglyceride or glycerophospholipid compound classes. Triglycerides are important for energy storage processes and are common biomarkers for assessing energy reserves (Conneely and Coates, 2023). The composition of glycerophospholipids in the lipid bilayer of cell membranes — phosphatidylethanolamine (PE), phosphatidylcholines (PC), phosphatidylserine (PS), phosphatidylinositol (PI), or phosphatidic acid (PA) — can shift due to temperature (van Meer et al., 2008), with previous research showing temperature-associated changes to PE and PC lipids in *C. maenas* (Chapelle, 1978; Chapelle, 1986).

### Integrative Analysis

Physiological and molecular datasets were integrated in two ways. First, a Weighted Correlation Network Analysis (WCNA) was used to understand correlations between compound abundance patterns, performance, and treatment conditions that significantly impacted both metabolomes and lipidomes using the `WGCNA` package (Langfelder and Horvath, 2008; Langfelder and Horvath, 2012). This analysis identified groups of co-expressed compounds, or module eigengenes, with similar abundance patterns, then correlated those eigengenes with righting response, oxygen consumption, and experimental conditions. Temperature was treated as a categorical variable to identify modules responding to multiple conditions (*ie.* generalized stress response). Module eigengenes significantly correlated (*P*-value < 0.05) with at least one experimental condition and physiological endpoint were considered for functional interpretation. Second, a Data Integration Analysis for Biomarker Discovery using Latent Variable Approaches for Omics Studies (DIABLO) (Rohart et al., 2017) was used to identify correlations between VIP metabolites and lipids. This supervised approach allows for the integration of metabolomic and lipidomic datasets in relation to experimental conditions. Prior to the DIABLO analysis, a sparse PLS (`pls`) was used to obtain an estimated correlation value between VIP metabolite and VIP lipid abundances. This estimated correlation was then used to determine the number of components that reduced classification error (`block.plsda`), and tune the metabolites and lipids to include in the analysis (`tune.block.splsda`). Finalized parameters were used to run the DIABLO analysis on VIP metabolites and lipids (`block.splsda`). A circos plot was used to visualize correlations between individual molecules with an absolute value greater than 0.70 (`circosPlot`).

## Results

### Performance Assays

#### Survivorship

Green crabs were held in experimental temperature conditions (ambient: 13.8°C ± 0.9°C; warm: 30.8°C ± 2.4°C; cold = 5.7°C ± 0.4°C ) for 22 days (**Figure S1**) with no meaningful difference in carapace width, weight, or sex ratios between the beginning and end of the experiment (**Figure S2**; additional information available in **Appendix 2**). Survival differed significantly across temperature treatments (**Figure 2**; Kaplan-Meier survival analysis: χ^2^_2_ = 18.7, *P*-value = 9 x 10^-5^). At 30°C, *C. maenas* had a 8.32-fold increase in mortality with each day of the experiment when compared with the 13°C control (Cox proportional hazards regression model (CPH): CI = 1.87-36.95, *P*-value = 0.005). There was no significant difference in hazard of death between crabs at 5°C and 13°C (CPH model: Hazard Ratio (HR) = 1.03, CI = 0.15-7.34, *P*-value = 0.97).

**Figure 2.**
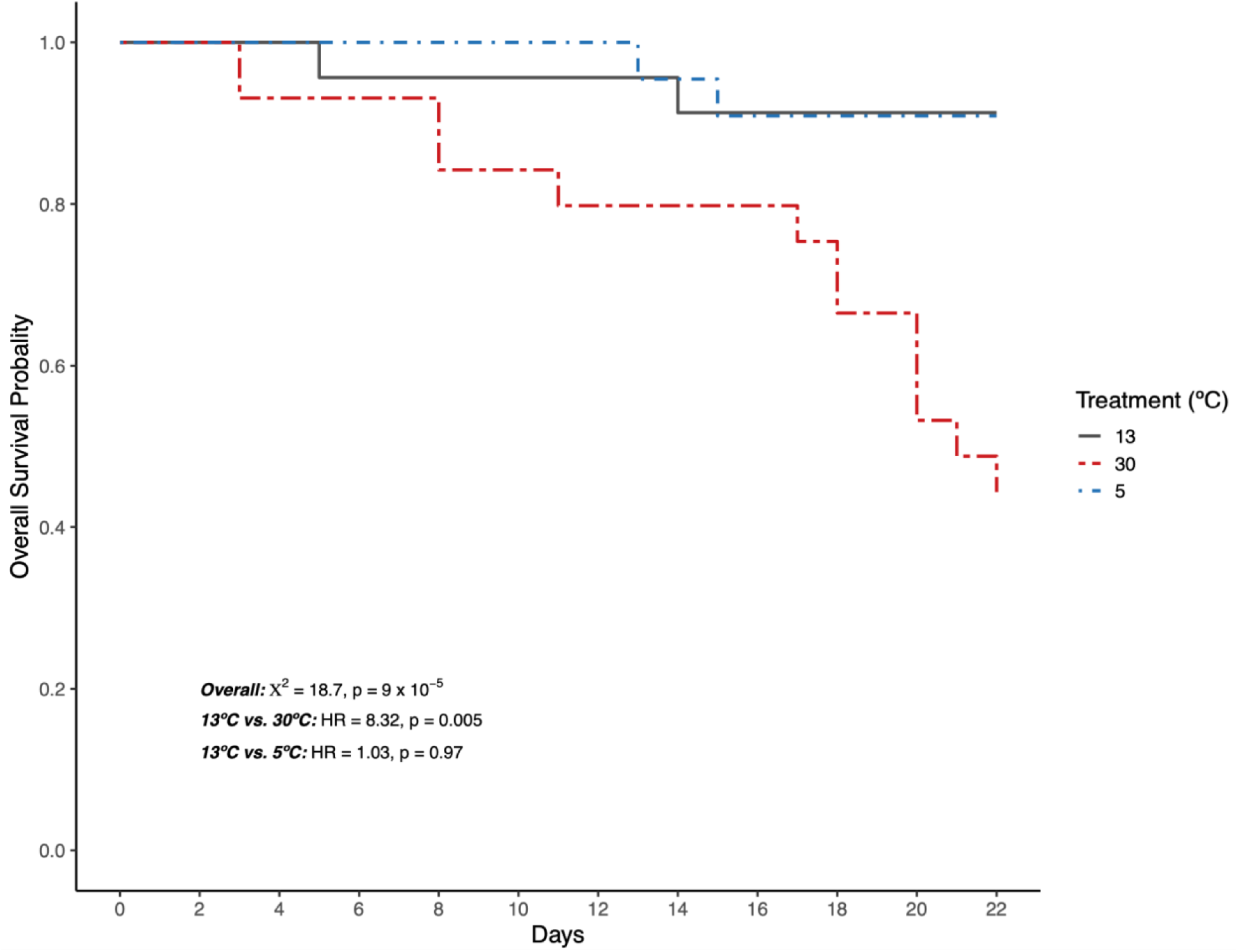
Proportion of surviving *C. maenas* throughout the experimental period. Grey, solid lines indicate survival of crabs at 13°C, while long, red dashes indicate survival at 30°C and short, blue dashes indicate survival at 5°C.

#### Time-to-right

Over the course of the experiment, there were five instances of crabs not righting within 90 seconds. One male crab at 5°C did not right on day 11, another male at 5°C did not right on day 15, and one male at 5°C, one male at 30°C, and one female at 30°C did not right on day 22. Four of these measurements were from crabs that did not right themselves during any trial, while the other measurement was from a crab that only righted during the first trial. These data points were excluded from further analysis.

TTR was significantly influenced by treatment (**Figure 3**). TTR for 5°C crabs was significantly higher than for 13°C (Tukey HSD_13°C vs. 5°C_: t = -15.48, *P*-value < 0.0001), while TTR for 30°C crabs was significantly lower than for 13°C crabs (Tukey HSD_13°C vs. 5°C_: t = 2.66, *P*-value = 0.02). The best fit model with the lowest AIC also included experimental day (GLM: t = -1.55, *P-*value = 0.12) and crab weight (GLM: t = 1.47, *P-*value = 0.14), although neither explanatory variable significantly impacted TTR.

**Figure 3.**
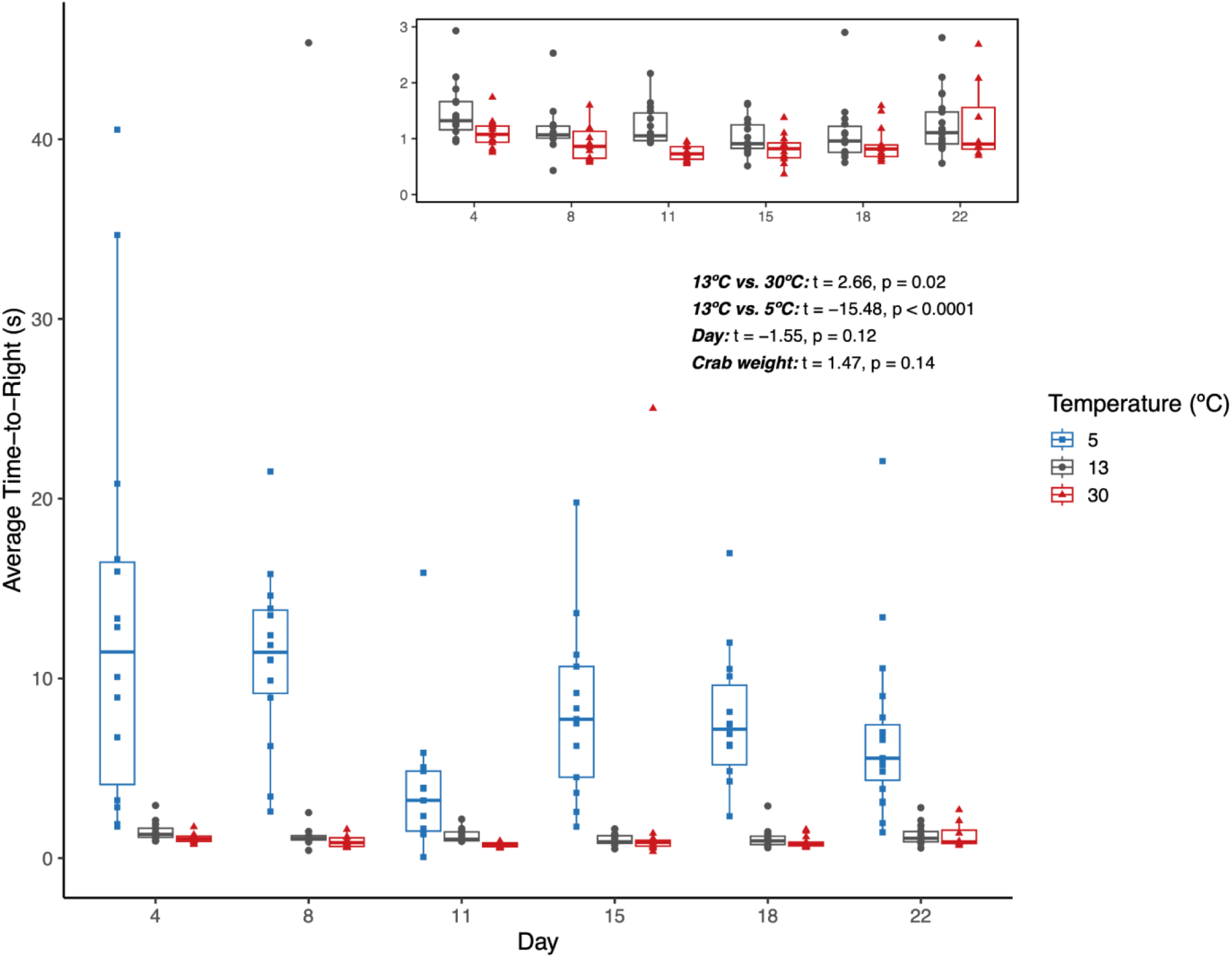
Average time-to-right (TTR) in seconds. Crabs in cold (5°C, blue squares), control (13°C, grey circles), and warm (30°C, red triangles) treatments are shown. The inset shows the data for the control and warm crabs between zero and three seconds. TTR was significantly influenced by treatment, with no significant influence of experimental day or the interaction between treatment and day.

#### Oxygen consumption

Temperature and day had a significant main effect on *C. maenas* oxygen consumption rate (**Figure 4**), with no impact of treatment x day interaction. Differences between temperature treatments were driven by significantly lower oxygen consumption at 5°C when compared to 13°C (Tukey HSD_13°C vs. 5°C_: t = 8.09, *P*-value < 0.0001). There was a marginal difference in oxygen consumption between 13°C and 30°C treatments (Tukey HSD_13°C vs. 30°C_: t = -2.38, *P*-value = 0.05). The significant influence of experimental day on oxygen consumption (GLM: t = 3.75, *P*-value < 0.0001) was driven by slower oxygen consumption at day 4 when compared to days 15 and 22 (**Table S1**).

**Figure 4.**
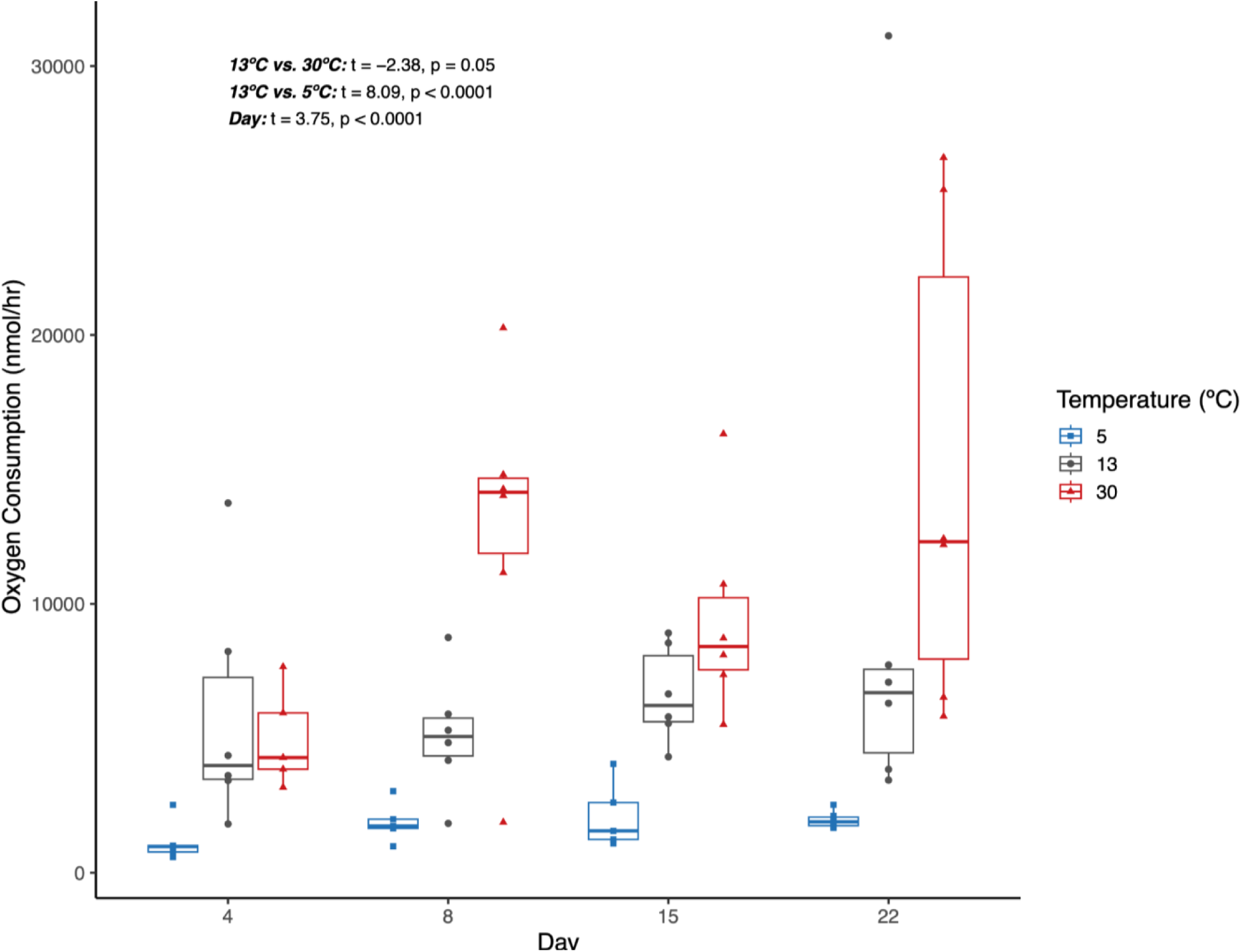
Oxygen consumption rate (nmol/hr) of *C. maenas*. Data for control (13°C, grey circles), warm (30°C, red triangles), and cold (5°C, blue squares) temperatures are shown at days 4, 8, 15, and 22. Temperature treatment and day significantly influenced oxygen consumption, with no impact of treatment x day interaction.

### Molecular Assays

#### Metabolomics

A total of 597 metabolites were identified through ALEX-CIS GCTOF MS. Of these, 152 metabolites were known compounds identified by the sequencer. A perMANOVA run on the known metabolites revealed significant effects of temperature (F = 3.08, *P*-value = 0.001), day (F = 1.97, *P*-value = 0.004), and their interaction on metabolite abundance (F = 1.66, *P*-value = 0.008 (**Figure S3**). Significant differences for all factors were due to centroid and dispersion differences (F_temperature_ = 5.58, *P*-value_temperature_ = 0.006; F_day_ = 6.04, *P*-value_day_ = 0.02; F_interaction_ = 4.90, *P*-value_interaction_ = 0.0008). A subsequent PLS-DA for known metabolites showed separation of *C. maenas* samples by treatment and day (**Figure 5A**).

**Figure 5.**
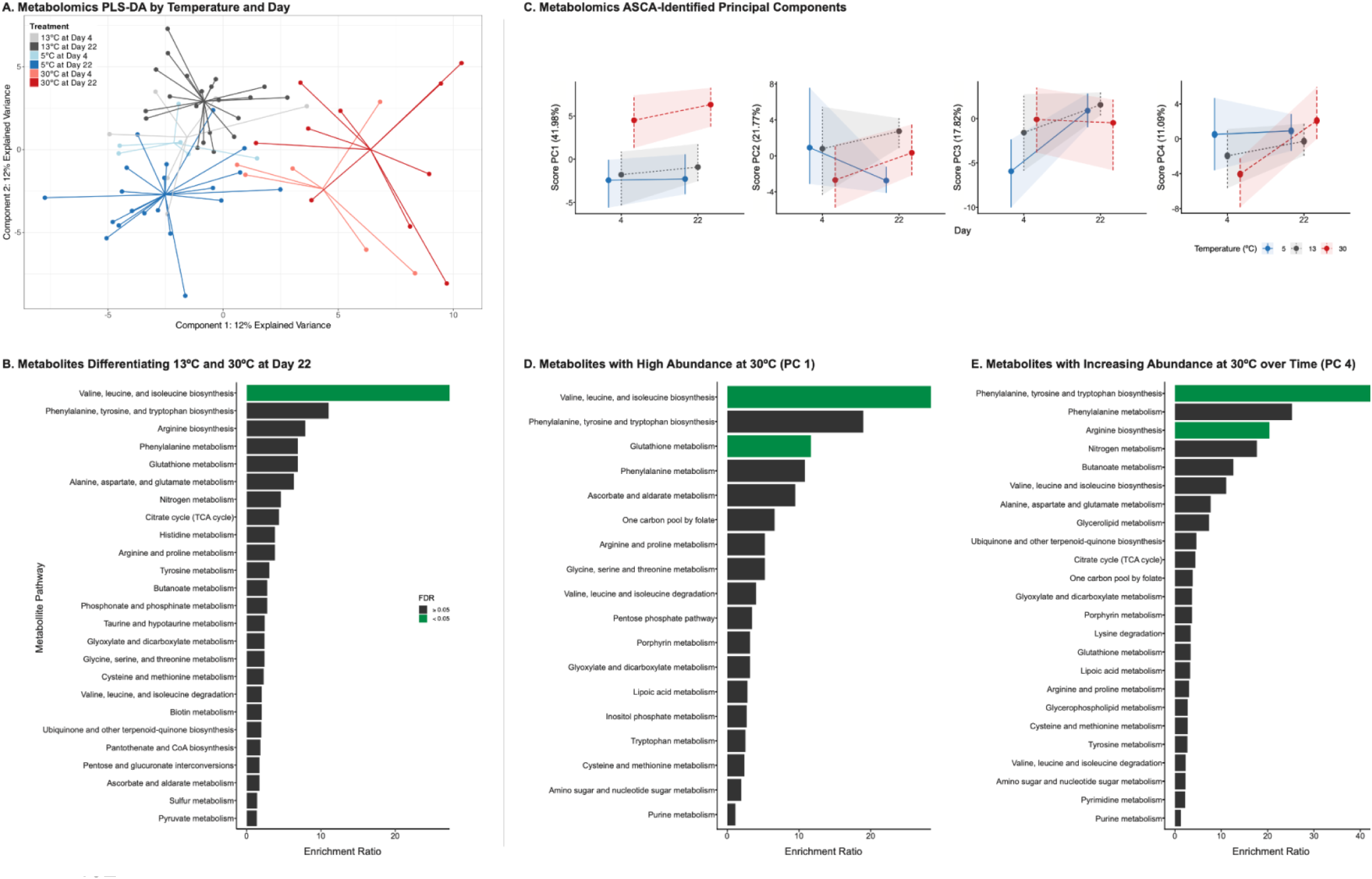
Metabolomics results. **A**) PLS-DA of log-transformed abundances for 152 known metabolites. Points represent abundances for individual crabs, with lines connecting each sample to the centroid for its respective treatment condition. **B**) Enrichment ratios (observed hits/expected hits) for 29 pairwise VIP metabolites driving differences between 13°C and 30°C metabolomes at the end of the experiment. **C**) Abundance patterns of metabolites organized into ASCA-identified Principal Components (PCs). Each PC explained > 10% of the total variance. Enrichment ratios for **D**) metabolites with high abundance throughout the experiment at 30°C (PC 1) and **E**) metabolites that were stable at 5°C but had increased abundance over time at 30°C (PC 4). For panels **B**), **D**), and **E**), pathways were identified using the Pathway Analysis module in the MetaboAnalyst v.6.0 interface, and significantly enriched terms (FDR < 0.05) are represented by green bars.

Metabolites with VIP scores > 1 and significant differences in log-normalized abundance (“pairwise VIP”) were identified to determine which compounds were most important for driving metabolome differences between pairs of temperature x day contrasts. While there were no pairwise VIP identified between 13°C and either 30°C or 5°C at day 4, there were pairwise VIP between temperature treatments at day 22 (VIP score ≥ 1 and adjusted *P*-value < 0.05). A total of 29 pairwise VIP differentiated the 13°C and 30°C treatments at the end of the experiment (**Table S2**). These VIP included metabolites important for respiration (malic acid; **Figure S4**) and calcium signaling (glutamine, glutathione, threonine, and tryptophan). Interestingly, there was no difference in the abundance of lactic acid, a biomarker for anaerobic respiration, between 13°C and 30°C at either timepoint (**Figure S4**). Pathway analysis demonstrated significant enrichment of the valine, leucine, and isoleucine biosynthesis pathway (FDR = 0.0004) (**Figure 5B**, **Table S3**). Differences between 5°C and 13°C at the end of the experiment were attributed to 31 metabolites (**Table S2**). These pairwise VIP were involved in valine, leucine, and isoleucine biosynthesis and arginine biosynthesis; however, these pathways were not significantly enriched (**Table S3**). Several of these VIP were also involved in cold protection (proline and alanine), calcium signaling (UDP-N-acetylglucosamine, serine, and threonine), and cellular respiration (alpha-ketoglutarate; **Figure S4**). Six pairwise VIP were identified when interrogating differences in metabolite abundance at 5°C between the beginning and end of the experiment, including citric acid cycle compound fumaric acid (**Table S2)**. Pairwise VIP metabolites were also screened for between 30°C and 5°C to understand if acclimation to temperature extremes employs unique metabolite pathways. The majority of 42 metabolites that differentiated 30°C from 5°C at the end of the experiment were also important for differentiating 13°C from either 30°C or 5°C at the end of the experiment (see **Appendix 3**).

An ASCA model identified four abundance patterns, or PCs, that each explained > 10% of the variance in metabolite abundance across temperature and day (**Figure 5C**). Metabolites positively loaded onto PC 1 had consistently higher abundances at 30°C over time (**Figure 5C**, **Figure S5A**). Pathway analysis of the top 20 metabolites correlated with PC 1 demonstrated significant enrichment of the valine, leucine, and isoleucine biosynthesis (FDR = 0.008) and glutathione metabolism (FDR = 0.009) pathways in these metabolites (**Figure 5D**). Metabolites positively loaded onto PC 2 decreased in abundance over time at 5°C, but increased in abundance over time at 30°C (**Figure 5C**, **Figure S5B**). PC 3 describes metabolites where positively loaded metabolites increase in abundance over time at 5°C (**Figure 5C**, **Figure S5C**).

There were no significantly enriched pathways represented by the top 20 compounds in PCs 2 or 3. Finally, PC 4 describes metabolites with relatively stable abundance at 5°C that also exhibited a more change in abundance at 30°C over time (**Figure 5C**, **Figure S5D**). Arginine biosynthesis (FDR = 0.02) and phenylalanine, tyrosine, and tryptophan biosynthesis (FDR = 0.03) were significantly enriched pathways for the top 20 metabolites in PC 4 (**Figure 5E**).

#### Lipidomics

Tandem MS/MS methods provided spectral data for 2,406 lipids across all samples. A total of 415 compounds had associated structural and/or nomenclature information, and were classified as “known” lipids. Known lipid profiles significantly differed by temperature (perMANOVA: F = 4.99, *P*-value = 0.001), and this significant difference was attributed to both centroid and dispersion differences (F = 4.54, *P*-value = 0.01) (**Figure S6**). Known lipid profiles also differed significantly by day (F_day,known_ = 2.74, *P*-value_day,known_ = 0.01), but this was not the case for the full set of detected compounds (F_day,all_ = 1.95, *P*-value_day,all_ = 0.07). There was no significant impact of temperature x day interaction on known lipid abundance (perMANOVA: F = 1.28, *P*-value = 0.18). Given that there is no biological distinction between “unknown” and “known” compounds and the known compounds represented < 20% of all detected compounds, we employed a conservative approach and only evaluated temperature effects in downstream analyses. A PLS-DA showed clear separation of lipids by temperature (**Figure 6A**).

**Figure 6.**
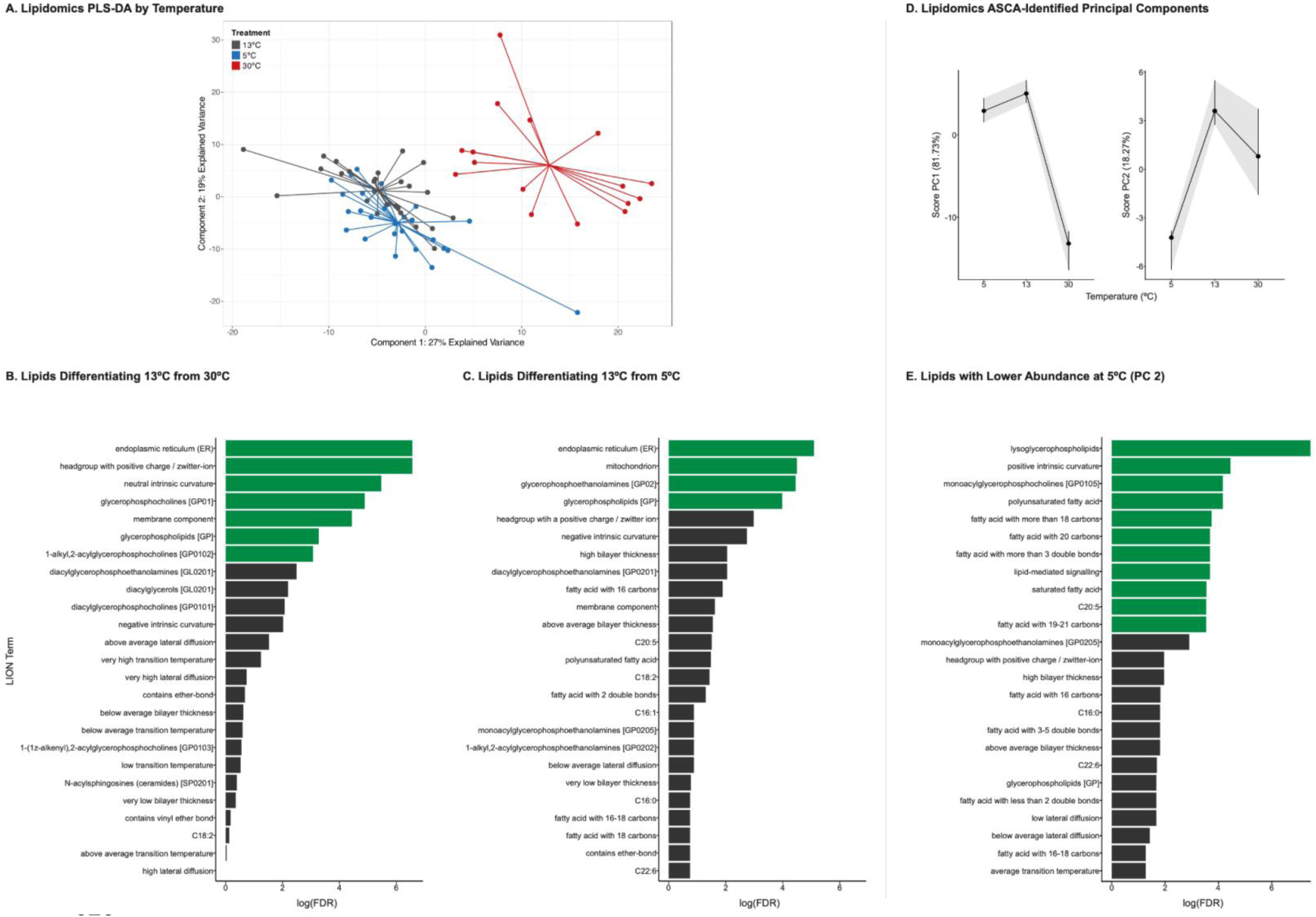
Lipidomics results. **A**) PLS-DA of log-transformed abundances for 415 known lipids. Points represent abundances for individual crabs, with lines connecting each sample to the centroid for its respective treatment condition. LION term enrichment results for **B**) 130 pairwise VIP lipids associated with differences between 13°C and 30°C and **C**) 61 pairwise VIP lipids associated with differences between 13°C and 5°C. **D**) Abundance patterns of lipids organized into ASCA-identified Principal Components (PCs). Each PC explained > 10% of the total variance. **E**) LION term enrichment results for lipids with lower abundance at 5°C (PC 2). For panels **B**), **C**), and **E**), enrichment analysis was conducted using LION/web target-list mode, and significantly enriched terms (FDR < 0.05) are represented by green bars.

The majority of lipids detected were glycerophospholipids, with PC and PE lipids highly represented as pairwise VIP lipids. A total of 130 lipids contributed to separation between 13°C and 30°C (VIP scores ≥ 1 and adjusted *P*-value < 0.05) (**Table S4**), with saturated and monounsaturated glycerophospholipids more abundant at 30°C. Enriched LION terms associated with these pairwise VIP lipids were endoplasmic reticulum (FDR = 0.001), headgroup with positive charge/zwitterion (FDR = 0.001), neutral intrinsic curvature (FDR = 0.004), glycerophosphocholines (FDR = 0.007), membrane component (FDR = 0.01), and glycerophospholipids (FDR = 0.04) (**Figure 6B**; **Table S5**). There were 61 lipids driving variation between 13°C and 5°C (**Table S4**). Only one PC lipid, PC 14:0_16:1, had a higher abundance in 5°C while all other glycerophospholipid VIPs had higher abundance in 13°C. Other pairwise VIP lipids with higher abundance in 5°C included polyunsaturated monoacylglycerophosphocholines (LPC) and monoacylglycerophosphoethanolamines (LPE) (**Table S4**). Enriched LION terms associated with pairwise VIP lipids were endoplasmic reticulum (FDR = 0.006), mitochondrion (FDR = 0.01), glycerophosphoethanolamines (FDR = 0.01), and glycerophospholipids (FDR = 0.02) (**Figure 6C**; **Table S5**). The majority of the 135 lipids driving differences between 30°C and 5°C were also important for distinguishing each temperature condition from the 13°C control (see **Appendix 3**).

An ASCA model identified two PCs that explained > 10% of total variance in lipid abundance across temperature (**Figure 6D**). PC 1 described lipids with lower abundance at 30°C when compared to 5°C and 13°C (**Figure 6D**, **Figure S7A**). There were no enriched LION terms represented in this PC. PC 2 described lipids with lower abundance at 5°C when compared to 30°C and 13°C (**Figure 6D**, **Figure S7B**). There were 11 enriched LION terms in this PC, including lysoglycerophospholipids (FDR = 0.0006), LPC (FDR = 0.02), and lipid-mediated signaling (FDR = 0.03) (**Figure 6E**).

### Integrative Analysis

There were three metabolites whose abundance was correlated with lipid abundance (**Figure 7A**, **Table S2**). Myo-inositol was negatively correlated with 11 lipids, which were predominantly unsaturated glycerophospholipids (**Figure S8**). Hydroquinone was negatively correlated with three lipids, all of which were also positively correlated with myo-inositol (**Figure S8**). Threonine abundance was positively correlated with the abundance of 11 lipids (**Figure S8**). These lipids were a mix of unsaturated cerines and unsaturated PC glycerophospholipids. Threnonine is also involved in the valine, leucine, and isoleucine biosynthesis pathway, which was enriched in the 30°C treatment at the end of the experiment.

**Figure 7.**
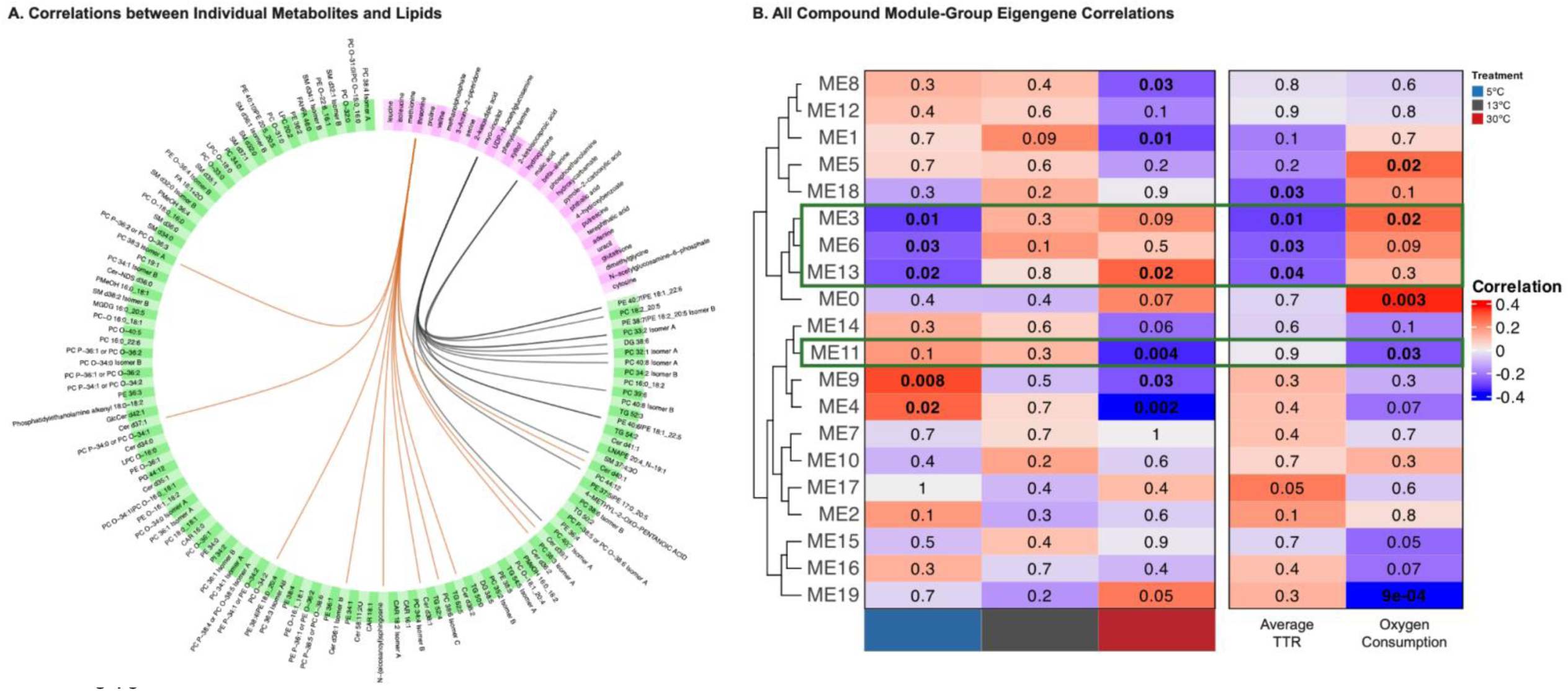
Integrative analysis results. **A**) Circos plot displaying correlations with absolute values above 0.7 between VIP metabolites (pink) and VIP lipids (green). Positive correlations are shown in brown while negative correlations are shown in black. **B**) Module-Group eigengene correlations from WCNA analysis. Temperatures are clustered by abundance similarity. Module eigengenes represent groups of compounds (known metabolites and/or lipids) that are co-expressed. Colors indicate the correlation between a module eigengene and either temperature, righting response, or oxygen consumption, with red colors representing positive correlations and blue colors representing negative correlations. *P*-values for each correlation are included, with bold values indicating significance (*P*-value < 0.05). Modules that were significantly correlated with temperature as well as righting response and/or oxygen consumption are outlined in green.

There were four modules of compounds significantly correlated with both temperature and at least one physiological trait (**Figure 7B**). Only one of these module eigengenes, ME 3, contained both metabolites and lipids. Compound abundance in ME 3 was negatively correlated with 5°C, negatively correlated with average TTR, and positively correlated with oxygen consumption (**Figure S9**). Compounds in this module include myo-inositol and several unsaturated glycerophospholipids. Abundance of the 21 lipids in ME 6 was negatively correlated with 5°C and righting response (**Figure S10**). These compounds were predominantly sphingomyelin lipids. The 11 lipids in ME 11 had abundances negatively correlated with both 30°C and oxygen consumption (**Figure S11**). These lipids were predominantly unsaturated. Abundance of the nine metabolites in ME 13 was negatively correlated with 5°C, positively correlated with 30°C, and negatively correlated with righting response (**Figure S12**).

## Discussion

Our results demonstrate the impacts of chronic thermal stress on crab performance and the importance of the metabolome and lipidome in driving physiological plasticity in response to temperature changes. As expected, crabs at 5°C exhibited consistently low metabolic activity (i.e., slower righting response and decreased oxygen consumption) while those at 30°C exhibited high metabolic activity (i.e., faster righting response and increased oxygen consumption). Interestingly, only crabs at 30°C had altered performance in line with our initial hypothesis, maintaining righting response and oxygen consumption early on, but having more variable oxygen consumption and higher mortality towards the end of the experiment. Specific molecular pathways enabled maintenance of motor activity and oxygen consumption at each temperature. Contrary to our expectations, there was no impact of time on the lipidome: alterations to the lipidome were consistently present after four days of exposure. These changes, in line with homeoviscous adaptation, may have facilitated motor activity across temperature. Metabolic reprogramming was ongoing throughout the thermal exposure, with significant differences in the metabolome at the beginning and end of the experiment. At each temperature, catabolism of various compounds provided energy necessary for aerobic respiration. These results highlight the central role of energy metabolism pathways in underpinning whole-organism physiological responses to temperature, and provide insight into how a highly invasive marine crab is able to thrive across a wide thermal range.

### Homeoviscous adaptation and related processes facilitate motor activity across temperatures

Righting response patterns highlight the plasticity of green crab thermal physiology. At all time points, crabs at 5°C exhibited a slower righting response than counterparts at 30°C, consistent with our understanding of ectotherm biology and prior research in the species (Jost et al., 2012; Rivers et al., 2025; Somero et al., 2017; Venkataraman et al., 2025). Previous acute temperature studies have shown that *C. maenas* righting response at thermal extremes is dynamic between 24-96 hours (Jost et al., 2012; Venkataraman et al., 2025). The lack of a significant impact of time on righting response in our study suggests that righting response acclimation occurs within four days, and that this acclimatory response can be sustained over several weeks.

We hypothesize that homeoviscous adaptation may have modified cell membrane structure to help maintain motor activity and righting response across temperatures. Cell and organelle membrane fluidity is dependent on temperature: high temperatures can liquify membrane structure and cold temperatures can increase membrane rigidity (Cossins and Prosser, 1978). To retain ideal membrane fluidity, organisms employ homeoviscous adaptation by regulating the abundance of saturated (no double bonds, associated with rigid membranes) and unsaturated (at least one double bond, associated with fluid membranes) glycerophospholipids in the cell membrane (Cossins and Prosser, 1978; Parrish, 2013; Sinensky, 1974). We observed higher abundances of saturated and monounsaturated glycerophospholipids at 30°C and lower abundances of these same compounds at 5°C. A higher proportion of saturated lipids at higher temperatures reduces membrane fluidity deeper in the bilayer (Cuculescu et al., 1995), providing structural integrity necessary for muscle contraction (Cossins and Prosser, 1978; Parrish, 2013; Sinensky, 1974). Conversely, replacing saturated and monounsaturated lipids with polyunsaturated lipids at 5°C may increase membrane fluidity to retain motor function in colder conditions. Similar patterns in glycerophospholipid abundance have been shown in the edible crab *Cancer pagurus* (Cuculescu et al., 1995), Atlantic hard clam *Mercenaria mercenaria* (Parent et al., 2008), and sea scallop *Placopecten magellanicus* (Hall et al., 2002), demonstrating core biological functions shared among marine invertebrates exposed to variable temperature environments.

Our results align with previous investigations of homeoviscous adaptation in *C. maenas*. Muscle tissue from male crabs had increased abundance of saturated and monounsaturated fatty acids and decreased abundance in polyunsaturated lipids after nine days of exposure to 27°C, while crabs exposed to 7°C displayed the opposite abundance patterns (Chapelle, 1978). These changes were observed after three days of thermal exposure, but were more pronounced after nine days. It is possible that Chapelle (1978) observed a significant impact of time due to the pairwise analytical approach they employed to examine abundance differences of 19 lipids, as opposed to our global lipidome analysis of 415 lipids. In both Chapelle (1978) and our study, differential abundance of PC and PE lipids were the main drivers of these changes, with limited changes observed in other glycerophospholipids or triglycerides (Chapelle, 1978). This suggests that PC and PE lipids are suitable biomarkers for exploring homeoviscous adaptation in future *C. maenas* or other marine invertebrate thermal acclimation studies.

Combined analysis of the metabolome and lipidome demonstrate the complexities of homeoviscous adaptation and motor activity regulation. We identified a module eigengene with nine metabolites whose abundance was positively correlated with temperature and negatively correlated with righting response time (ME 13). Three of these compounds were saturated fatty acids, which are regulated through homeoviscous adaptation to maintain membrane structure. Saturated fatty acids like stearic acid (Ali and Szabó, 2023; Hac-Wydro et al., 2009), arachidic acid (precursor to arachidonic acid), and heptadecanoic acid (Brash, 2001; Liang et al., 2004), had higher abundance at high temperature, likely contributing to increased membrane rigidity. Conversely, these fatty acids had lower abundances at low temperatures, increasing membrane fluidity. While regulation of membrane structure itself can indirectly facilitate motor function, other compounds involved in muscle contraction and signaling processes are more directly connected to motor function maintenance. Two polyamines found in ME 13, putrescine and spermidine, are known to enable muscle contraction and regulation in several crustaceans (Elgavish et al., 1984; Lucena et al., 2017), and in mammals (Fernández et al., 1995; Galasso et al., 2023; Pegg, 2016). Higher abundance of these compounds at 30°C suggests that they played a role in increased motor function, and therefore faster righting response at high temperature. Similarly, lower polyamine abundance at 5°C may contribute to slower righting responses in cold conditions. Another module eigenene, ME 6, was negatively correlated with both 5°C and righting response. The majority of the lipids in this module were sphingomyelins (SM) with lower abundance at 5°C. In addition to being minor cell membrane components, SM are crucial for cell signaling and membrane trafficking (Chakraborty and Jiang, 2013; Fabri et al., 2020). Low abundances of SM appears to be important for organisms that regularly experience colder temperatures. For example, SM(Ivanisevic et al., 2011)cold-adapted *Arigiope bruennichi* spiders experiencing cold temperatures (Ortiz-Movliav et al., 2024)(Ivanisevic et al., 2011) The low abundance of these molecules in crabs at 5°C may have resulted in slower cellular processes, which in turn could slow righting response. (Ivanisevic et al., 2011)(Chapelle, 1978)(Ortiz-Movliav et al., 2024)

### Metabolic reprogramming to meet energetic demands of thermal acclimation

#### High temperature elicits catabolism of branched-chain amino acids

Our investigation of thermal physiology across multiple weeks elucidates potential mechanisms of metabolic compensation green crabs employ when facing chronic thermal stress. An increase in environmental temperature increases metabolic rates in ectotherms, which in turn increases energy demand (Pörtner, 2001; Pörtner, 2002; Pörtner, 2010; Somero et al., 2017). According to the prevailing framework for crustacean thermal tolerance, the oxygen- and capacity-limitation of thermal tolerance hypothesis (OCLTT), as organisms transition out of their optimal thermal range into the pejus range, they employ metabolic compensation to meet energetic demands (Pörtner, 2001; Pörtner, 2002; Pörtner, 2010). While the increased oxygen consumption at high temperatures reported in this and previous studies (McGaw and Whiteley, 2024; Nancollas and McGaw, 2021; Rodrigues et al., 2015) is indicative of *C. maenas* attempting to meet energetic demand, organisms in the pejus range are typically unable to meet oxygen demand by changing respiration rates alone. Organisms may start to employ anaerobic respiration, leading to increased abundance of anaerobic metabolites (Pörtner, 2001; Pörtner, 2002; Pörtner, 2010; Verberk et al., 2016). Interestingly, *C. maenas* at 30°C did not have significantly different lactic acid abundance than counterparts at 13°C or 5°C, suggesting that anaerobic metabolism was not employed more during chronic high temperature exposure.

Previous research has focused on acute mechanisms that may allow *C. maenas* to temporarily extend its thermal optimum range without using anaerobic metabolism (Giomi and Pörtner, 2013; Weber et al., 2008), but it is unlikely that these mechanisms can be maintained long-term. Taken together, our data suggest that crabs are employing alternative mechanisms that facilitate aerobic respiration in the pejus range under chronic exposure.

Given that anaerobic metabolism was likely not sustained by crabs under chronic heat stress, crabs may have used metabolic reprogramming to meet the energetic demands of acclimatization and homoviscous adaptation. Metabolic reprogramming comprises the regulation of core metabolic functions to maintain homeostasis across a range of conditions (Moore et al., 2025; Wang et al., 2022). The catabolism of other metabolites can provide additional energy to supplement classic aerobic metabolism in response to thermal stress. At elevated temperature, we observed significant enrichment of valine, leucine, and isoleucine biosynthesis, suggesting that crabs utilized energy from catabolism, or degradation, of branched-chain amino acids (BCAAs). The abundance of BCAAs can regulate metabolic processes such as glucose intake, lipid metabolism, lipogenesis, and gluconeogenesis in crustaceans (Xie et al., 2023). Crustaceans derive BCAAs from their diet as they are unable to synthesize them (Neinast et al., 2019). Therefore, it is likely that the enrichment of “biosynthesis” identified by MetaboAnalyst is in fact related to increased catabolism of BCAAs at higher temperatures. High temperature can upregulate branched-chain aminotransferases, which in turn degrade BCAA abundance (Xu et al., 2017). Degradation of BCAAs then generates carbons that can then be used for aerobic metabolism in the citric acid cycle (Neinast et al., 2019). The low abundances of BCAAs at 30°C suggests that crabs required continual catabolism of BCAAs to meet higher energetic demand (**Table S2**). Similar processes have been observed in other marine invertebrates exposed to short-term stressors. For example, low BCAA abundance has been implicated as a marker for increased energy demand to maintain homeostasis in shrimp exposed to high ammonia (Xiao et al., 2019), and the sea cucumber *Apostichopus japonicus* regulates BCAA catabolism after 48 hours of exposure to elevated temperature (Xu et al., 2016; Xu et al., 2017). The enrichment of BCAA catabolism after 22 days of high temperature exposure suggests that this catabolic process can be maintained as long as BCAAs are available in the diet and catabolism is not energetically costly. Since researchers have been unable to identify the transition from aerobic to anaerobic metabolism in *C. maenas* using traditional physiological metrics (Giomi and Pörtner, 2013; Jost et al., 2012), future work should interrogate the use of BCAAs as biomarkers using both gene expression and metabolomics to identify critical thermal thresholds.

#### Compensatory mechanisms complement metabolic depression exhibited at cold temperatures

Crabs exposed to chronic cold temperature conditions in our study exhibited mechanistic responses consistent across arthropod species: metabolic depression (Klein Breteler, 1975; Rivers, 2024), amino acid and protein catabolism (Colson-Proch et al., 2010; Zhu et al., 2024), and use of cryoprotectant molecules ((Churchill and Storey, 1989; Dou et al., 2019; Shang et al., 2015). Metabolic depression allows organisms to match metabolic demand with lower energy intake as a result of reduced feeding or activity at lower temperatures (Pörtner, 2010). Consistent with observations from previous research, crabs at 5°C maintained a low level of oxygen consumption consistent with metabolic depression, which may have allowed them to withstand chronic cold temperatures (Klein Breteler, 1975; Rivers, 2024).

Although metabolic depression would reduce metabolic demand, crabs at 5°C would still require energy to maintain minimal motor function. Similar to counterparts in elevated temperatures, crabs in cold temperatures may have employed amino acid and protein catabolism for energy production. Arginine biosynthesis was significantly enriched in metabolites that had stable, high abundance in cold conditions. Given that arginine is important for detoxification of ammonia waste produced by amino acid and protein catabolism (Colson-Proch et al., 2010), it is possible that crabs were engaging in catabolic processes and required arginine for waste detoxification. Additionally, arginine itself can be catabolized to produce ATP by marine invertebrates and cold-adapted insects (Colson-Proch et al., 2010; Zhu et al., 2024). While we did not detect any arginine in our study, we did detect higher abundances of intermediate compounds produced during arginine catabolism — proline and alanine — at 5°C. Additionally, we detected higher abundance of proline during early exposure, and higher abundance of alanine at the end of the experiment. These patterns suggest that *C. maenas* may have been producing arginine for immediate catabolism into proline followed by alanine for energy production to supplement energy acquired from reduced feeding activity. More direct measurement of protein catabolism in future studies is necessary to understand how this process can contribute to cold tolerance.

Presence of intermediate compounds like proline and alanine in *C. maenas* exposed to cold temperatures could also be indicative of cryoprotection mechanisms. Low molecular weight cryoprotectant molecules, such as osmolytes and by-products of anaerobic metabolism, are important for lowering the freezing point of the internal environment, minimizing cell dehydration, and protecting cell membranes and proteins from damage (Gill et al., 2024; Lee, 2010; Storey and Storey, 1996; Storey and Storey, 2017). While the specific mechanism of action is unclear, increased concentrations of these compounds have been detected in several invertebrates exposed to cold or freezing conditions. Proline is a known cryoprotectant in overwintering insects (Churchill and Storey, 1989; Dou et al., 2019; Shang et al., 2015), and proline supplementation in diets has been shown to extend cold tolerance of insects (Dou et al., 2019; Koštál et al., 2012) and marine crustaceans (Zhu et al., 2024). Both proline and alanine concentration increased in *Geukensia demissus* mussels acclimated to low temperature, suggesting they are important for cold tolerance (Murphy, 1977). Although crabs were not exposed to freezing temperatures in this experiment, higher abundance of these osmolytes could indicate a need to minimize cellular damage at lower temperatures. Research that can localize these molecules to specific regions of the cell can provide insight into their use for cold acclimation.

The potential of *C. maenas* to utilize alternative energy pathways and cryoprotectants may help to explain the impressive cold tolerance exhibited by this temperate species, which can survive below 5°C for at least four months (Kelley et al., 2013) and below 2°C for at least two months (Rivers et al., 2025). The species also thrives in sub-arctic conditions in its non-native range in Newfoundland at temperatures substantially below those it experiences in its native European range (Best et al., 2017). Future research should explore how long these compensatory mechanisms can be used for in order to determine their efficacy in enabling cold acclimation in this species, and potentially in predicting its continued spread in high latitudes (Kelley et al., 2013).

### Conclusion

Changes to ocean temperatures can elicit unprecedented species redistributions in both native and introduced species (Donelson et al., 2019; Fredston et al., 2021). Organisms that can maintain performance under chronic exposure to stressful temperatures will likely benefit from predicted temperature shifts, either by withstanding warmer ocean temperatures in their natural environment, or moving poleward into newly-accessible habitats (Donelson et al., 2019). Therefore, understanding the mechanisms employed during long-term application can provide insight into which species may be successful in new thermal regimes. Our work demonstrates how integrating methods across biological hierarchy can provide insights into the acclimatory responses used by organisms experiencing chronic thermal stress. By pairing metabolomics and lipidomics with performance assays for the first time in *C. maenas*, we show how metabolic reprogramming increases energy availability and maintains energy balance for cellular respiration, and how changes to membrane structure and small molecule abundance can underlie motor activity maintenance. Induction of these phenotypically plastic mechanisms may underlie the exceptionally flexible thermal physiology exhibited by *C. maenas* and its success as a marine invader. Similar methods can be used to examine how various other factors impact energy metabolism, including nutrient limitation (Beauclercq et al., 2023; Costa et al., 2024; Sogin et al., 2016), reproduction (Fu et al., 2022), and other environmental stressors (Venkataraman and Huffmyer, 2025). By holistically assessing the physiological plasticity underlying long-term acclimation, future work can make more informed predictions about species persistence or redistribution.

## Acknowledgements

We are grateful to Dr. Ariana S. Huffmyer for her insight into metabolomic and lipidomics analyses, and for reviewing draft text. We also thank Dr. Shelly Wanamaker for additional support on metabolomic and lipidomics methods and Dr. Christopher Murray for his assistance with respirometry methods.

## Data and Resource Availability

All raw data, scripts, results, and supplementary material are available at https://github.com/yaaminiv/green-crab-metabolomics.

## Appendices

## Appendix 1. Metabolomics sample preparation

Metabolite sample processing was performed at the UC Davis West Coast Metabolomics Center. Metabolites were extracted using the Matyash extraction procedure which includes MTBE, MeOH, and H_2_O. The bottom, aqueous phase was dried down and submitted to derivatization for gas chromatography. For derivatization, samples were first shaken at 30°C for 1.5 hours. Then 91 µL of MSTFA and FAMEs were added to each sample, and the samples were shaken again at 37°C for 30 minutes to finish derivatization. Samples were then vialed, capped, and injected onto the instrument. A 7890A GC coupled with a LECO-TOF was used to obtain metabolomics data. Derivatized sample (0.5 µL) was injected using a splitless method onto a RESTEK RTX-5SIL MS column with an Intergra-Guard at 275°C with a helium flow of 1 mL/min. The GC oven was set to hold at 50°C for one minute, then ramp up 20°C/min to reach 330°C, then finally hold at 330°C for five minutes. The transferline was set to 280°C while the EI ion source was set to 250°C. Data was collected from 85m/z to 500m/z at an acquisition rate of 17 spectra/sec.

## Appendix 2. Experimental design

Green crabs were held in experimental temperature conditions for 22 days after a seven-day acclimation period at 13°C. Experimental temperature conditions were 13.8°C ± 0.9°C, 30.8°C ± 2.4°C, and 5.7°C ± 0.4°C for ambient, warm, and cold treatments, respectively (**Figure S1**). While there were some temporary fluctuations with water exchanges, temperatures over the duration of the experimental period remained significantly different from each other (**Figure S1**; ANOVA: F = 42,002.94, *P*-value < 0.001).

Demographic information was assessed to confirm an even distribution of crabs by carapace width, weight, integument color, and sex throughout the experiment. At day 1 and day 22, there was no difference in carapace width and *C. maenas* weight (**Figure S2**) between temperature treatments. Integument color was significantly different between temperatures on day 22, likely due to an increased proportion of crabs with red-orange coloration at 30°C; this was expected, as color becomes more orange as crabs approach their next molt, and increased temperatures speed up intermolt intervals (Reid et al., 1997) (**Figure S2**; Chi-squared: *P*-value = 0.003). Sex ratio did not differ by temperature at either time point (**Figure S2**).

## Appendix 3. VIP identification between 30°C and 5°C

Pairwise VIP compounds were identified between 30°C and 5°C to understand how temperature extremes correspond with metabolome and lipidome shifts. Nine metabolites identified as pairwise VIP between 30°C and 5°C at day 4 had higher abundance at 30°C (**Table S2**). These metabolites were involved in the significantly enriched phenylalanine metabolism (FDR = 0.04), phenylalanine, tyrosine, and tryptophan biosynthesis (FDR = 0.02), and valine, leucine, and isoleucine biosynthesis (FDR = 0.04) pathways (**Table S3**). A total of 42 pairwise VIP were identified between 30°C and 5°C at day 22 **Table S2**), with ten metabolites having higher abundance in 30°C and the rest having higher abundance in 5°C. Of these pairwise VIP, 27 were also important for differentiating 13°C from either 30°C or 5°C at the end of the experiment. The 11 pairwise VIP metabolites unique to this contrast were not involved in any significantly enriched pathways (**Table S3**). A similar analysis was conducted to identify pairwise VIP lipids driving differences between 30°C and 5°C. A total of 135 pairwise VIP lipids were identified, 86 having higher abundance at 30°C and 9 having higher abundance at 5°C. A total of 28 lipids were uniquely important for driving differences between the two temperature extremes (**Table S4**). These lipids were not associated with any enriched LION terms (**Table S5**).

## Appendix 4. Supplementary Tables

**Table S1.** Pairwise statistical test results examining differences in righting responses between experimental days. Reported *P*-values were adjusted for multiple comparisons using the Bonferroni correction.

| Contrast | t | P-value |
| --- | --- | --- |
| Day 4 vs. Day 8 | -2.62 | 0.05 |
| Day 4 vs. Day 15 | -3.04 | 0.02 |
| Day 4 vs. Day 22 | -4.06 | $7.5 \times 10^{-4}$ |
| Day 8 vs. Day 15 | -0.42 | 0.97 |
| Day 8 vs. Day 22 | -1.44 | 0.48 |
| Day 15 vs. Day 22 | -1.02 | 0.74 |

**Table S2**. **Significant pairwise metabolite VIP identified for all contrasts.** https://github.com/yaaminiv/green-crab-metabolomics/blob/main/supplementary-material/metabolite-VIP-lists.xlsx

**Table S3. Pathway analysis results from MetaboAnalyst using pairwise metabolite VIP**. https://github.com/yaaminiv/green-crab-metabolomics/blob/main/supplementary-material/metabolite-enrichment-results.xlsx

**Table S4. Significant pairwise lipid VIP identified for all contrasts.** https://github.com/yaaminiv/green-crab-metabolomics/blob/main/supplementary-material/lipid-VIP-lists.xlsx

**Table S5. Enrichment analysis results from LION/web using pairwise lipid VIP**. https://github.com/yaaminiv/green-crab-metabolomics/blob/main/supplementary-material/lipid-enrichment-results.xlsx

**Table S6.** Metabolite and lipid correlations from supervised DIABLO analysis. Three metabolites were correlated with several lipids each. A 0.70 correlation cutoff was used.

| Metabolite | Negatively Correlated Lipids | Positively Correlated Lipids |
| --- | --- | --- |
| myo-inositol | PE 40:7\|PE 18:1_22:6,<br>PC 18:2_20:5<br>PC 33:2 Isomer A<br>DG 38:6<br>PC 32:1 Isomer A<br>PC 40:8 Isomer A<br>PC 39:6<br>PE 40:6\|PE 18:1_22:5<br>SM 37:4;3O<br>PC 44:12<br>PC 40:7 Isomer A | N/A |
| hydroquinone | PE 40:7\|PE 18:1_22:6<br>PC 33:2 Isomer A<br>PE 40:6\|PE 18:1_22:5 | N/A |
| threonine | N/A | PC 38:3 Isomer A<br>Cer d37:1<br>PC O-34:2<br>Cer d36:1 Isomer B<br>N-(eicosanoyl)sphingosine<br>Cer d38:1<br>PC 38:6 Isomer C<br>Cer d38:2<br>PC 36:3 Isomer A<br>Cer d39:1<br>Cer d40:1 |

## Appendix 5. Supplementary Figures

**Supplementary Figure 1.**
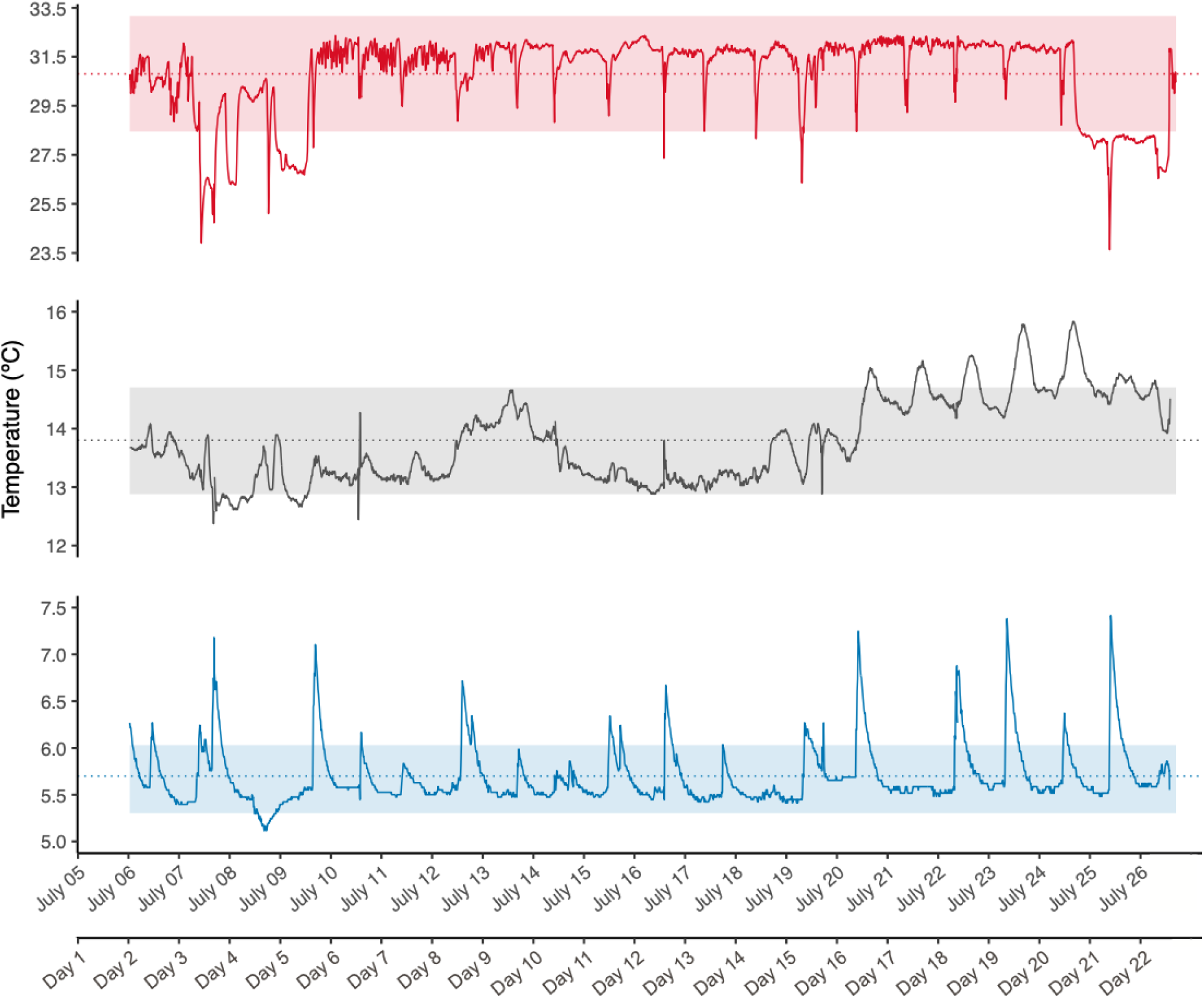
Temperatures over the course of the experimental period (July 5, 2022-July 26, 2022). The warm treatment is shown on top (30.8°C ± 2.4°C; red), control treatment in the middle (13.8°C ± 0.9°C; grey), and cold treatment on the bottom (5.7°C ± 0.4°C; blue). Average temperatures for each treatment are depicted by dashed lines, with standard error shown as shading. A secondary x-axis indicates experimental day (Day 1-Day 22).

**Supplementary Figure 2.**
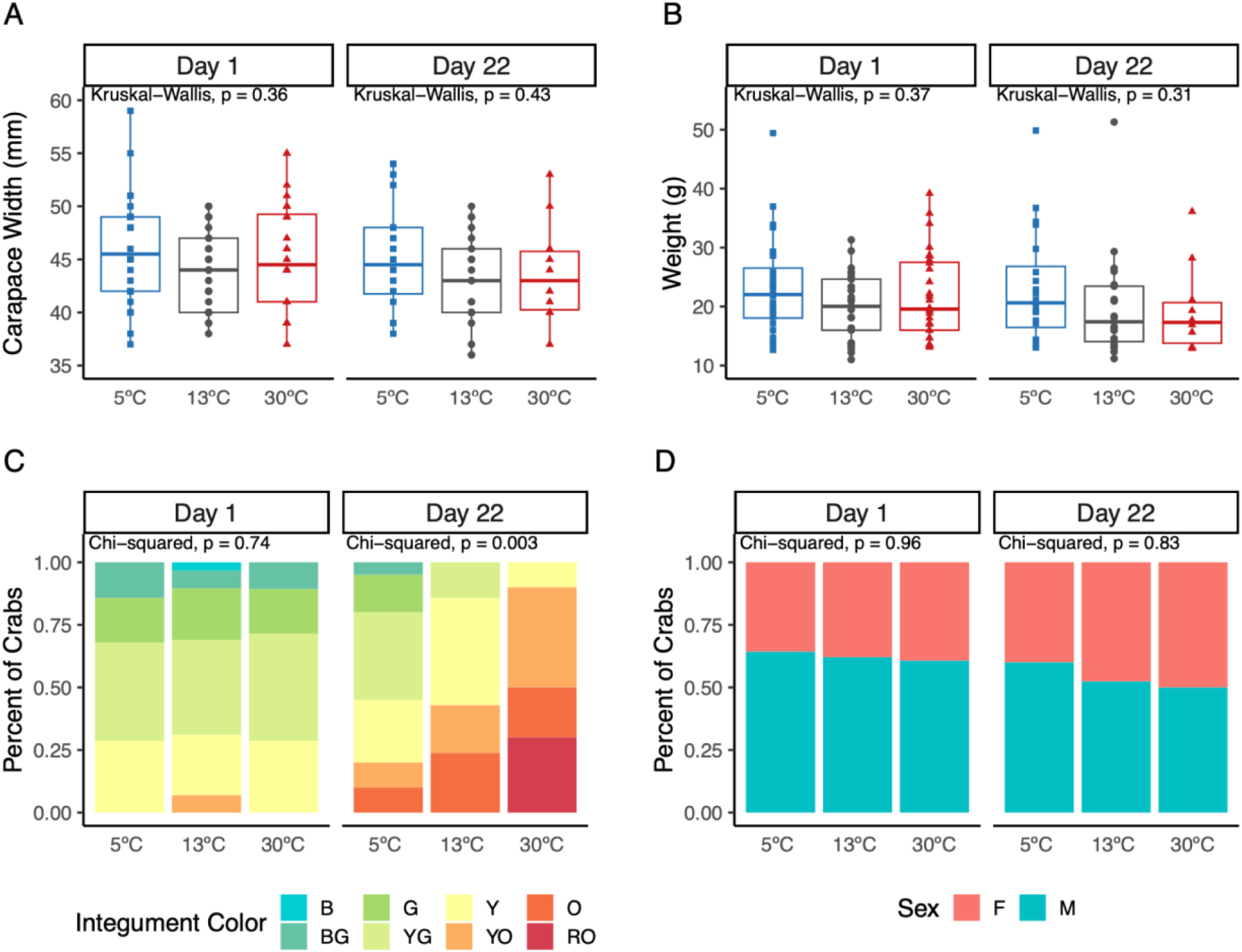
Demographic variables across 5°C, 13°C, and 30°C at the beginning (day 1) and end (day 22) of the experiment. For **A**) carapace width and **B**) crab weight measurements, mean values were compared between treatments at each time point using a Kruskal-Wallis test. Differences in the distribution of **C**) integument color (B: blue; BG: blue-green; G: green; YG: yellow-green; Y: yellow; YO: yellow-orange; O: orange, RO: red-orange) and **D**) sexes across treatments was assessed using chi-squared tests. On day 22, integument color was significantly different between the three temperatures. There was no difference in carapace width, crab weight, or sex ratio by temperature during the experiment.

**Supplementary Figure 3.**
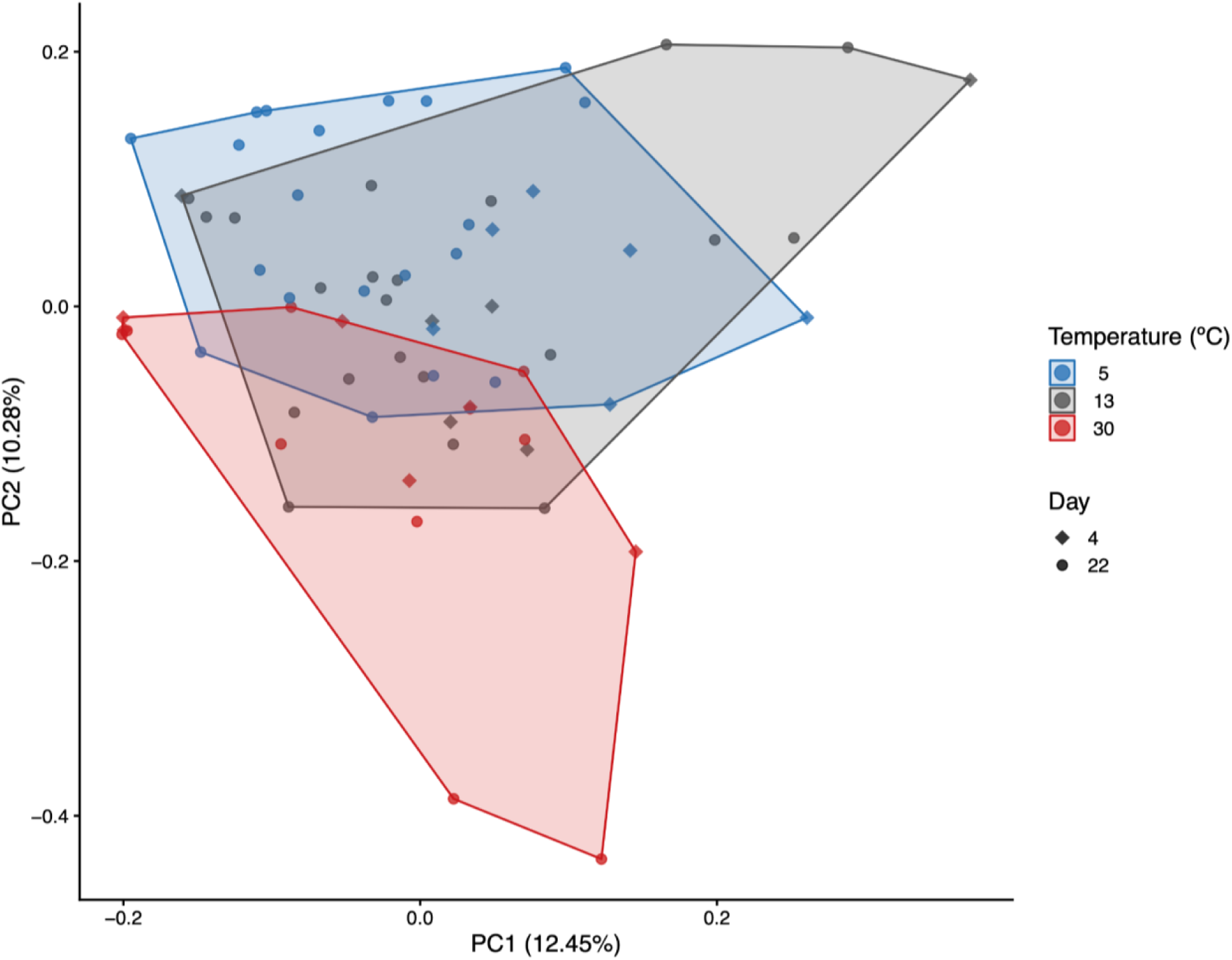
PCA for 152 known metabolites. Known metabolites are compounds which were identified by the sequencer. Colors distinguish between temperature treatments, while shape differentiates between experimental day. A perMANOVA identified significant differences between global metabolite abundance by temperature, day, and the interaction between temperature and day.

**Supplementary Figure 4.**
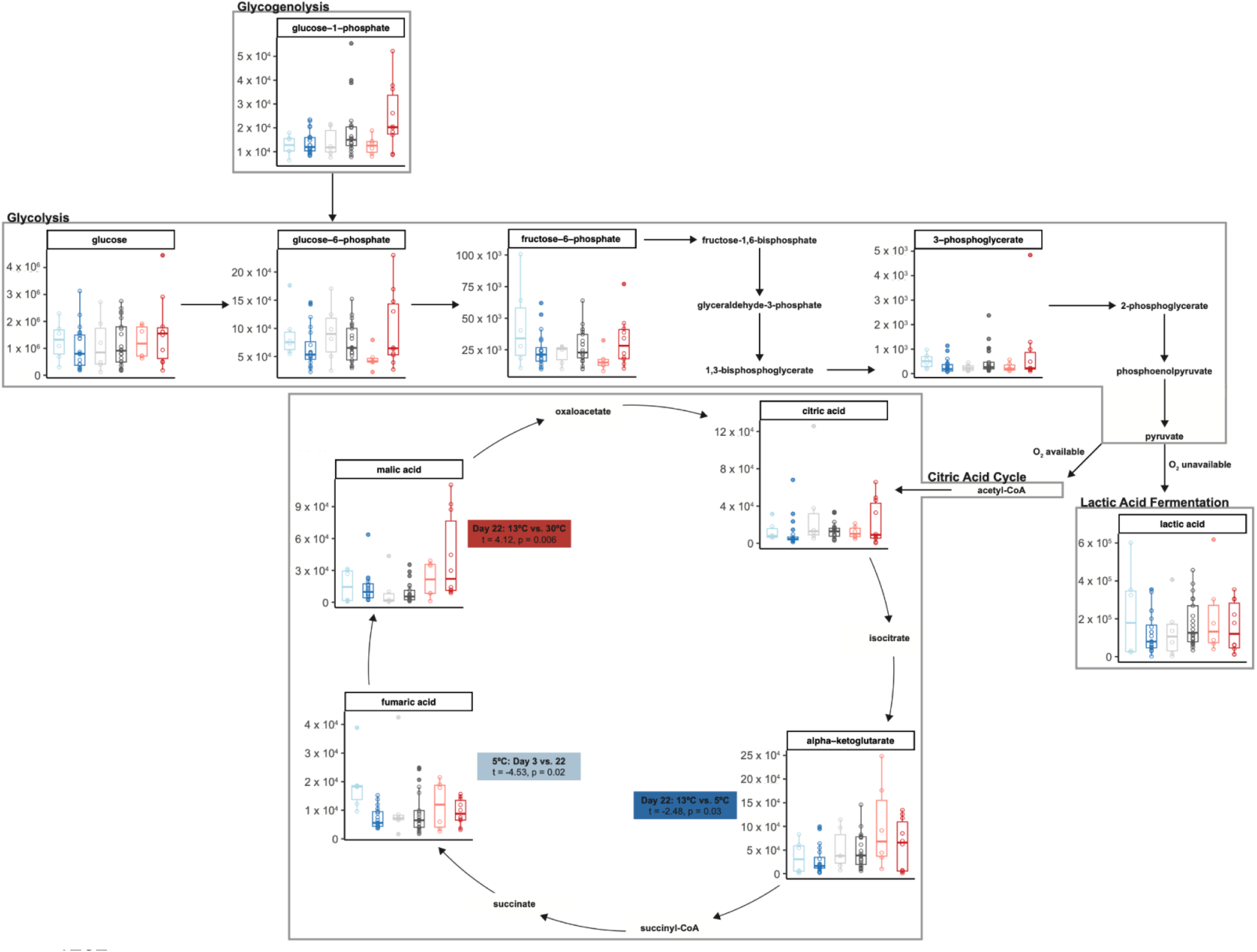
Comparison of abundance for metabolites involved in aerobic and anaerobic metabolism. Abundances are presented for detected metabolites involved in cellular respiration processes, with colors representing temperatures (blue = 5°C, gray = 13°C, red = 30°C) and shades representing day (lighter shades = day 4, darker shades = day 22). Three metabolites were significant pairwise VIP for different contrasts: alpha-ketoglutarate (Day 22: 13°C vs. 5°C), fumaric acid (5°C: Day 4 vs. 22), and malic acid (Day 22: 13°C vs. 30°C).

**Supplementary Figure 5.**
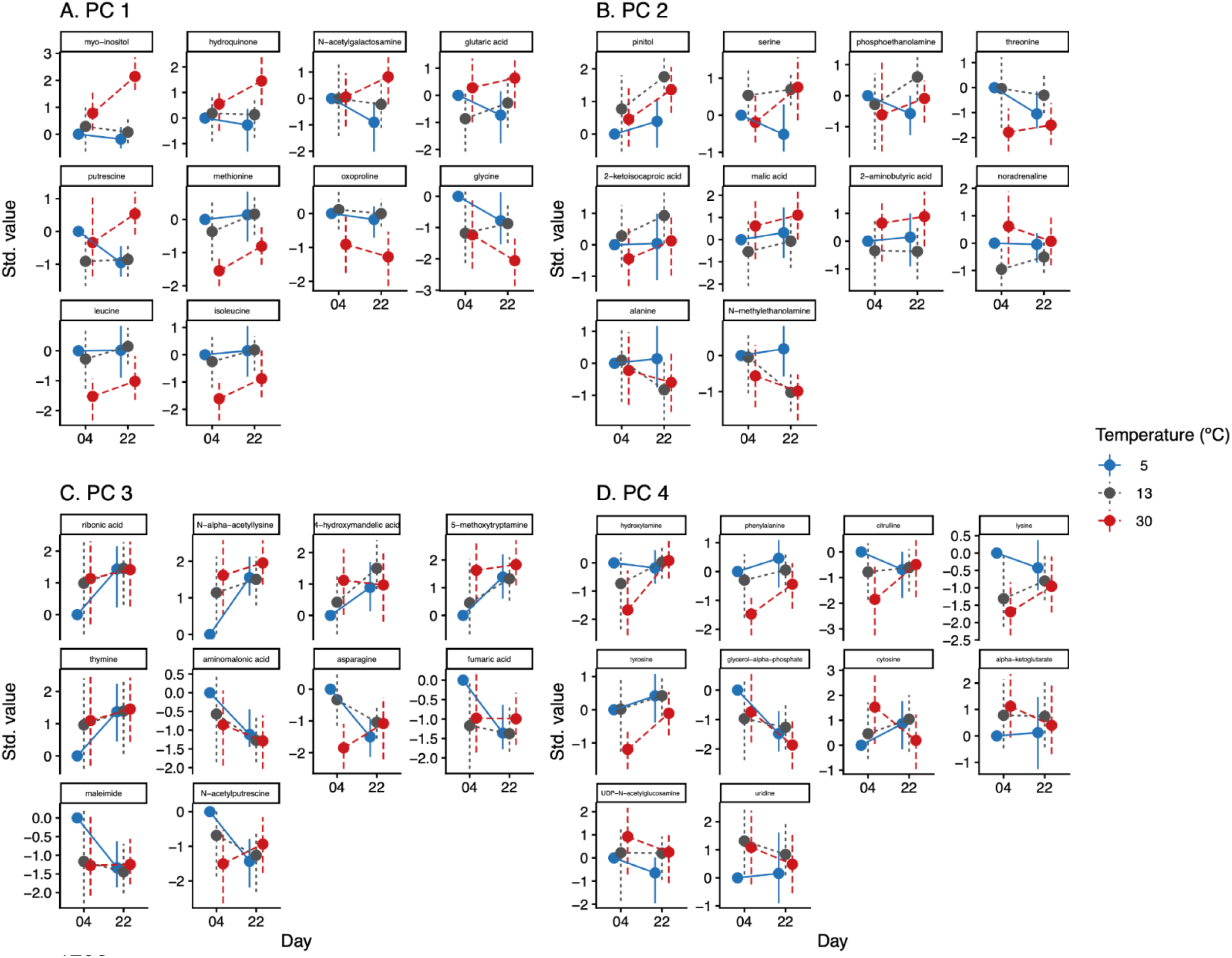
Marginal means for metabolites strongly loaded onto A) PC 1 (41.98%), B) PC 2 (21.77%), C) PC 3 (17.82%), and D) PC 4 (11.09%). For each PC, the top five metabolites with the strongest positive loadings and the top five metabolites with the strongest negative loadings are shown.

**Supplementary Figure 6.**
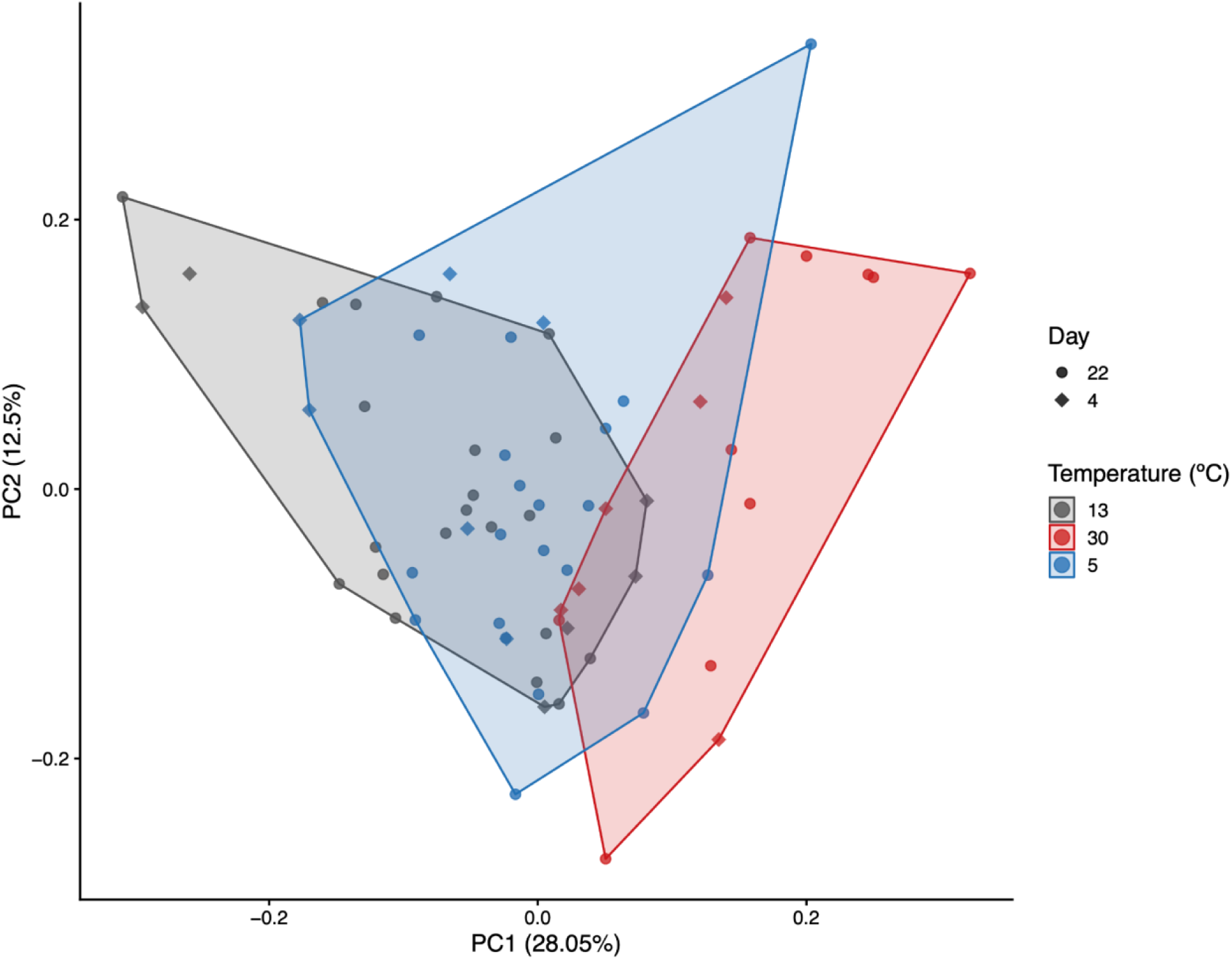
PCA for 415 known lipids. Known lipids are defined as those identified by the sequencer. Colors distinguish between temperature treatments, while shape differentiates between experimental day. A perMANOVA identified a significant impact of temperature on the global lipidome, with no significant impact of day of the interaction between temperature and day.

**Supplementary Figure 7.**
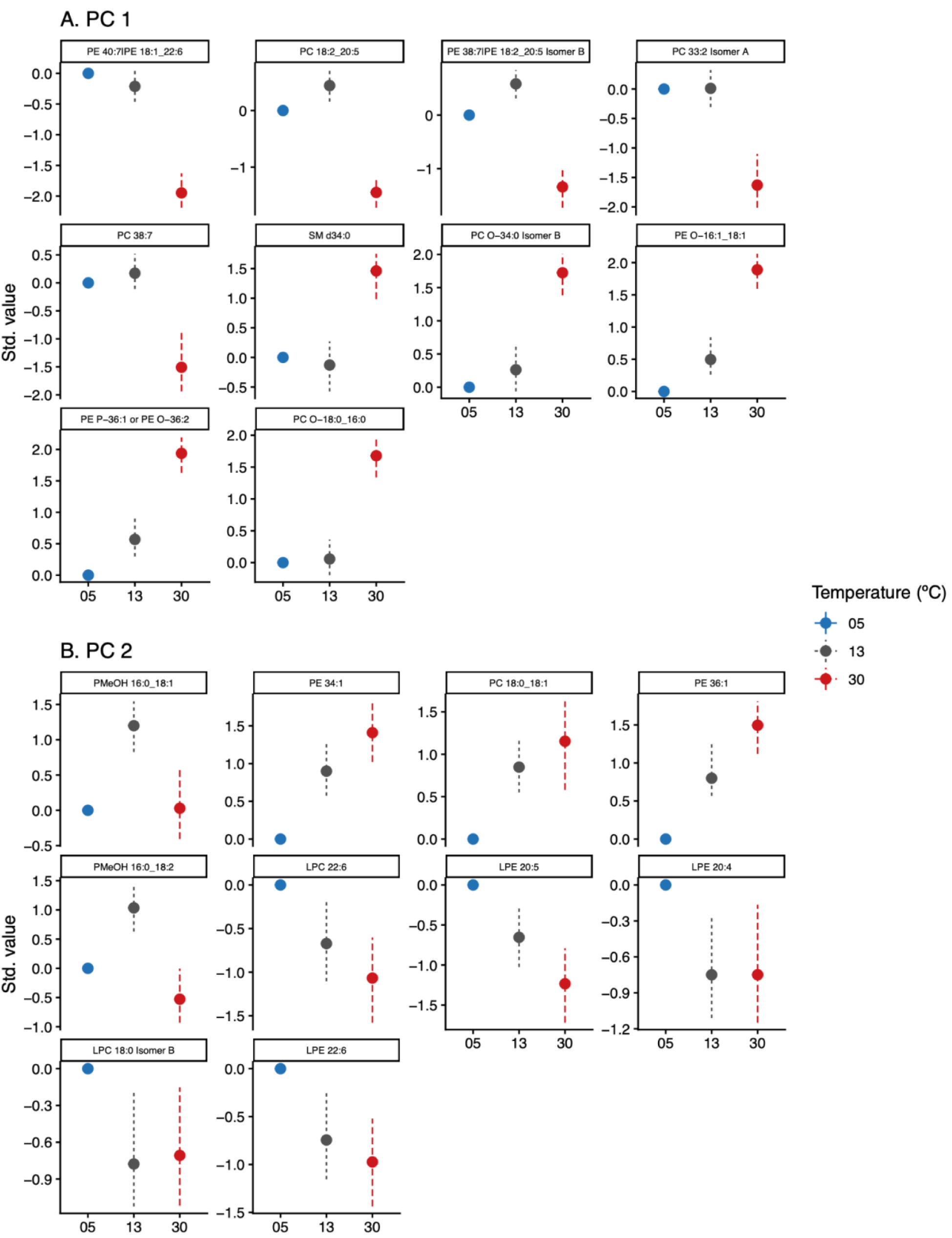
Marginal means for lipids strongly loaded onto A) PC 1 (81.73%) and B) PC 2 (18.27%). For each PC, the top five lipids with the strongest positive loadings and the top five lipids with the strongest negative loadings are shown.

**Supplementary Figure 8.**
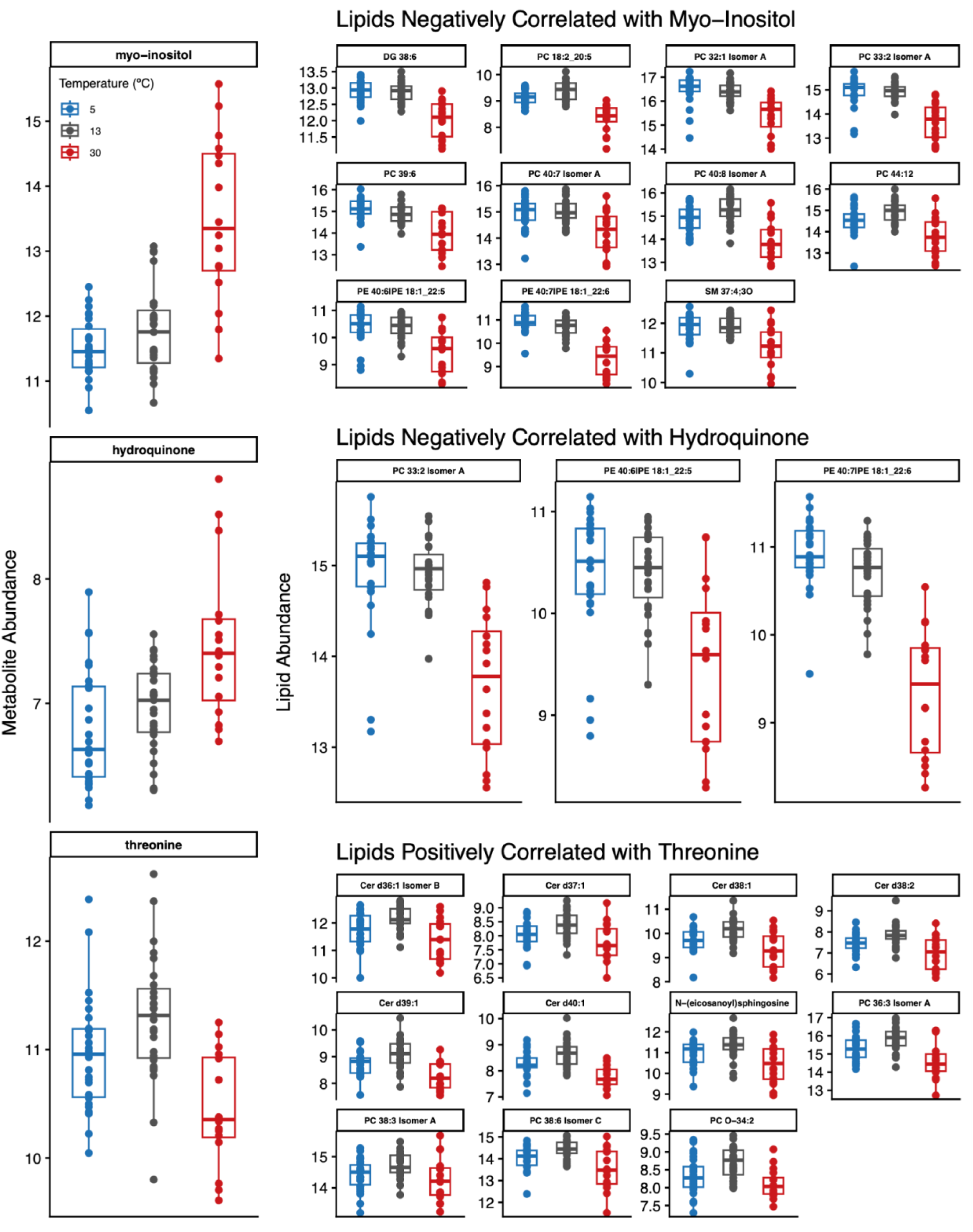
Abundance of strongly correlated (> 0.70) metabolites and lipids correlated (> 0.70) across temperature. A DIABLO analysis identified correlations between the metabolites myo-inositol, hydroquinone, and threonine and several lipids. Lipid abundance was either negatively correlated with myo-inositol and hydroquinone or positively associated with threonine. Several lipids were correlated with multiple metabolites.

**Supplementary Figure 9.**
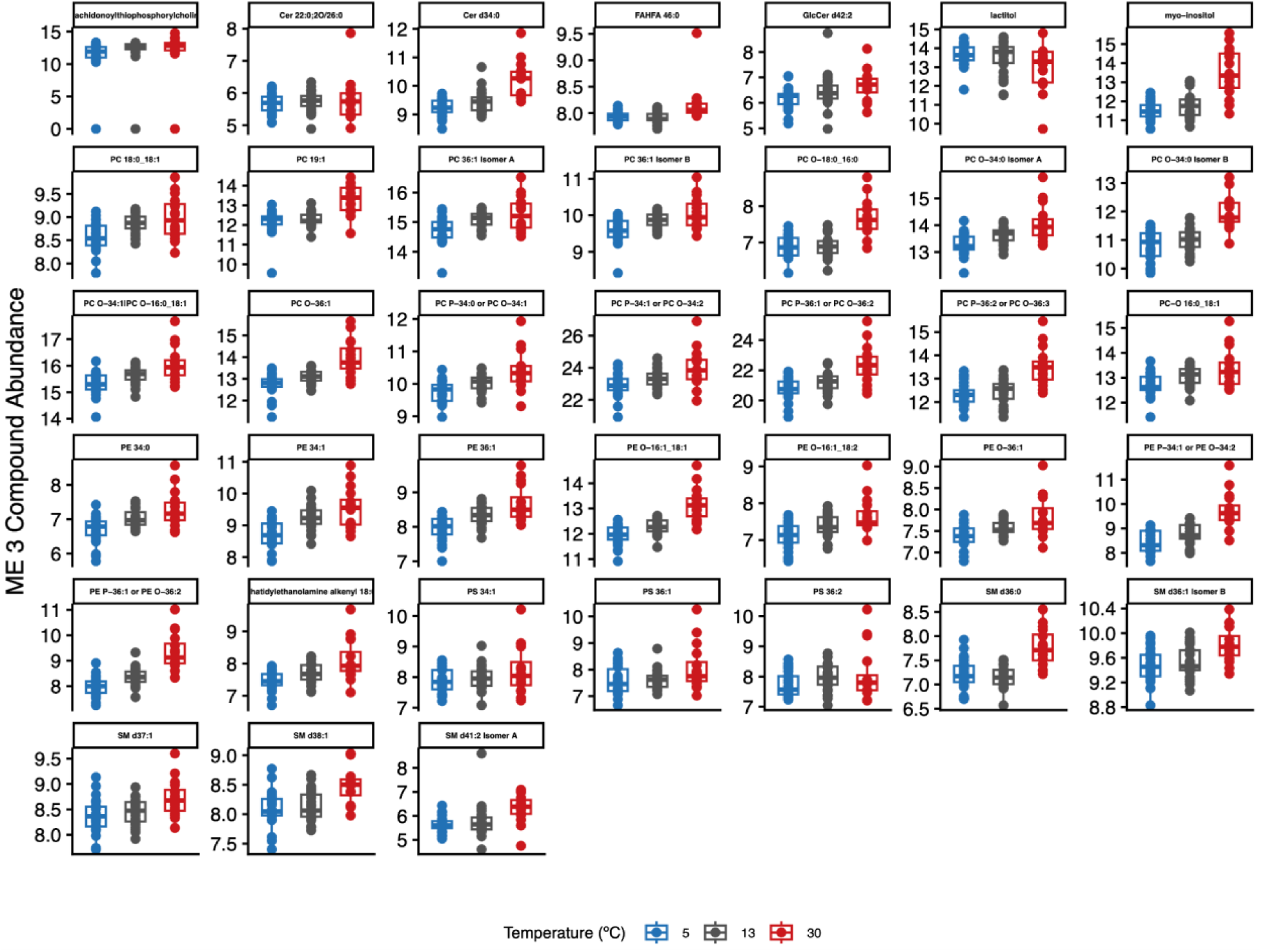
Abundance of 38 compounds (metabolites and lipids) in ME 3 across temperature. Compound abundance in ME 3 was negatively correlated with 5°C, negatively correlated with average TTR, and positively correlated with oxygen consumption.

**Supplementary Figure 10.**
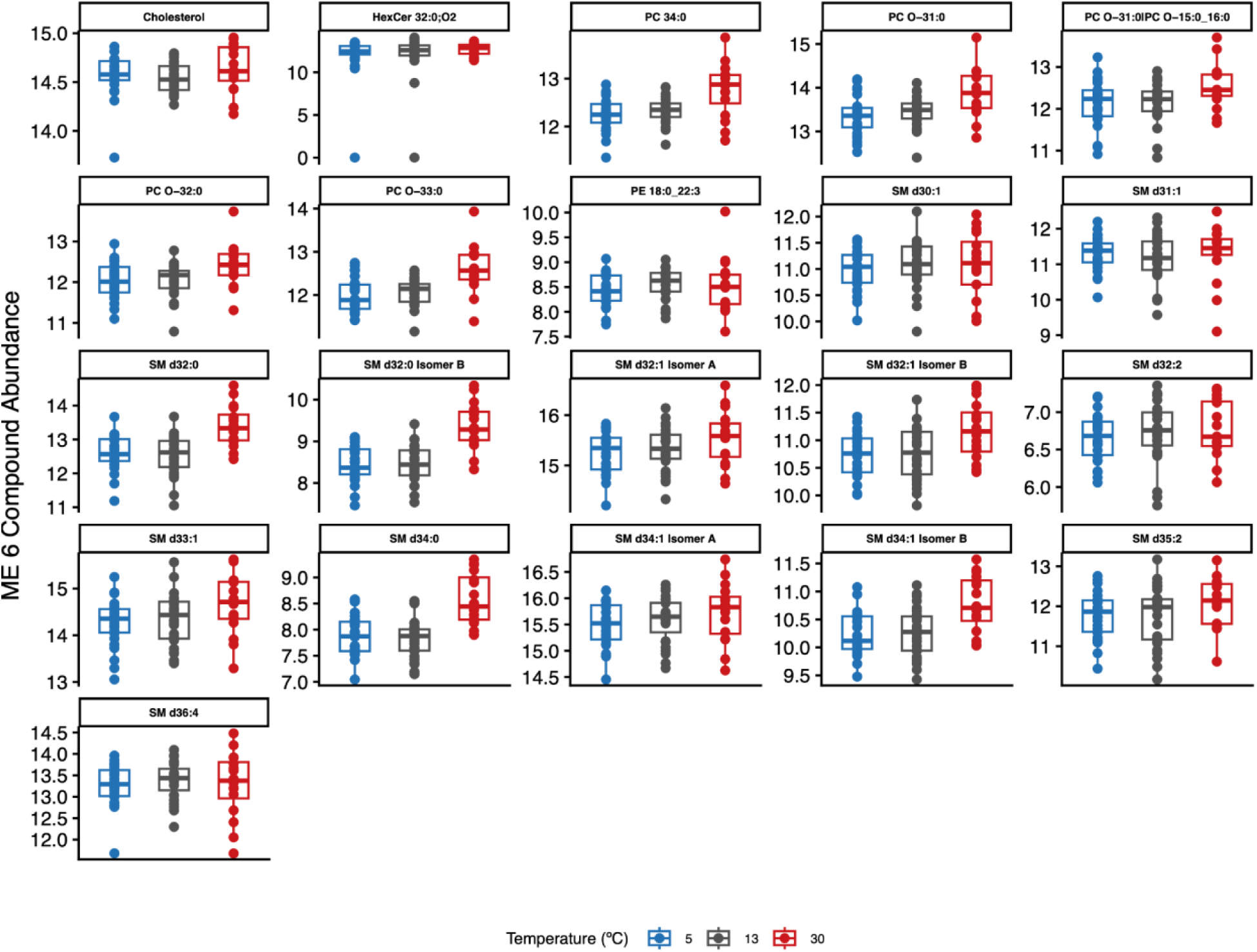
Abundance of 21 lipids in ME 6 across temperature. Lipid abundance in ME 6 was negatively correlated with 5°C and righting response.

**Supplementary Figure 11.**
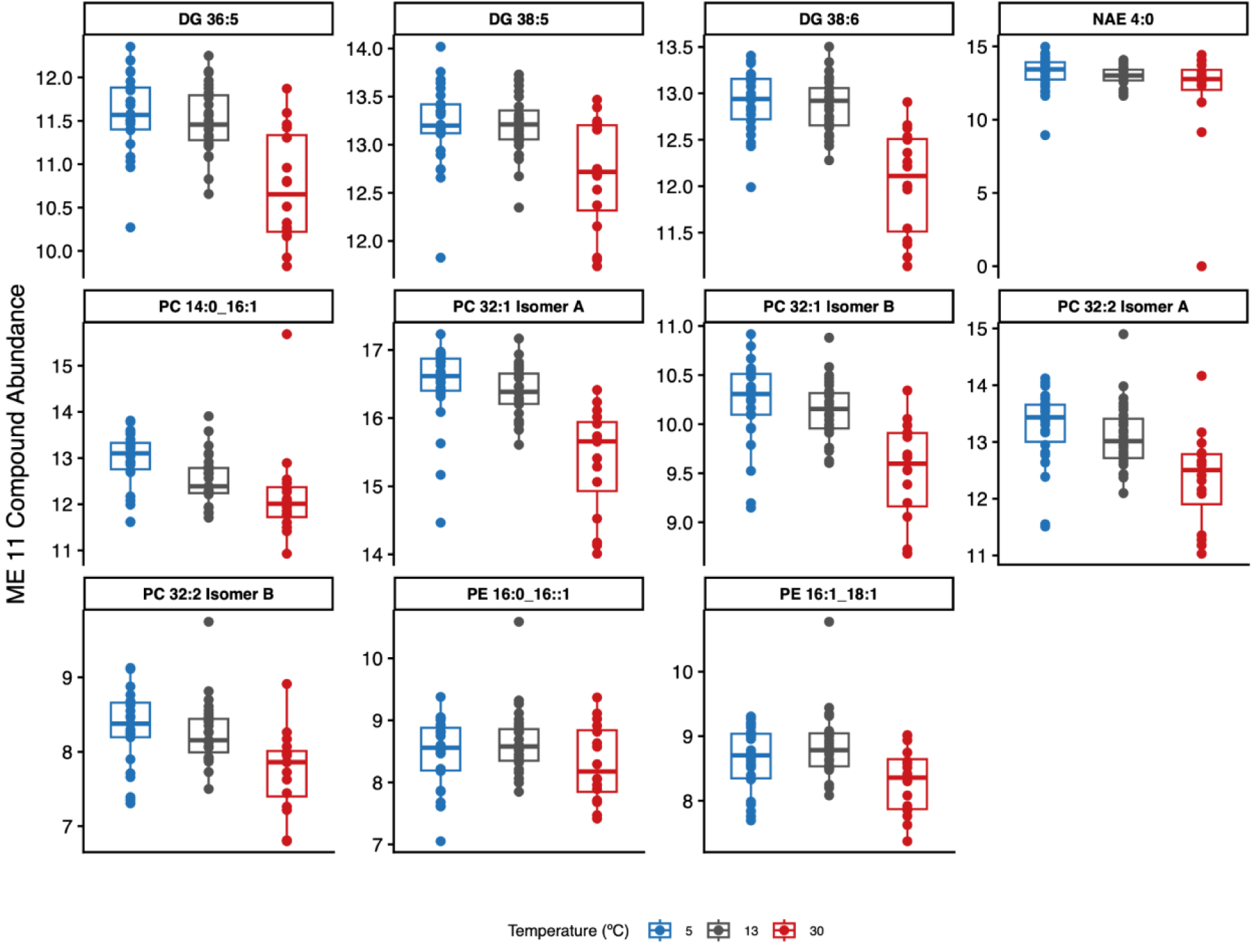
Abundance of 11 lipids in ME 11 across temperature. Lipid abundance in ME 11 was negatively correlated with 30°C and oxygen consumption.

**Supplementary Figure 12.**
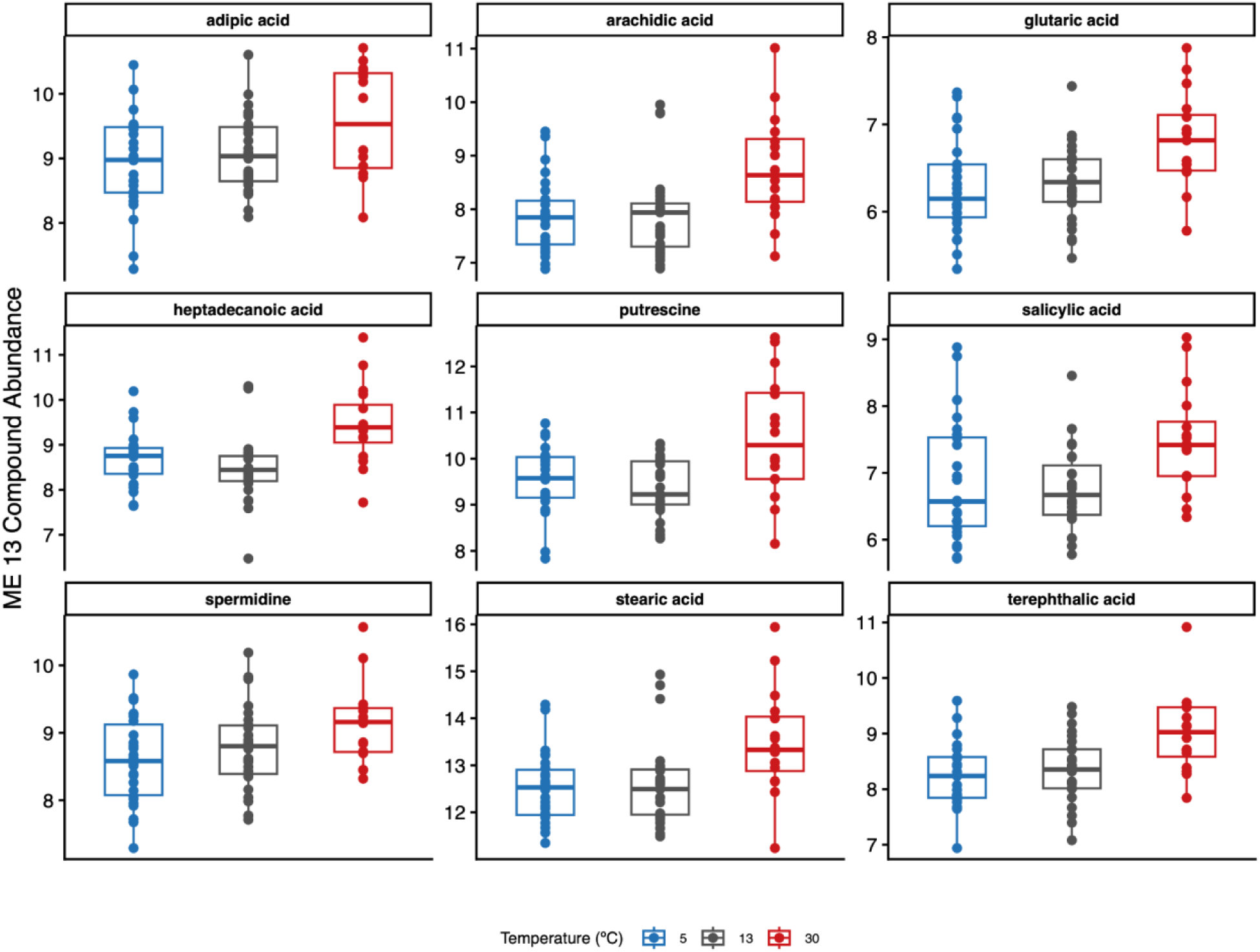
Abundance of nine metabolites in ME 13 across temperature. Metabolite abundance in ME 13 was negatively correlated with 5°C, positively correlated with 30°C, and negatively correlated with righting response.

